# A RNF12-USP26 amplification loop promotes germ cell specification and is disrupted in urogenital disorders

**DOI:** 10.1101/2020.11.16.378398

**Authors:** Anna Segarra-Fas, Francisco Bustos, Rachel Toth, Gino Nardocci, Greg M. Findlay

**Affiliations:** The MRC Protein Phosphorylation and Ubiquitylation Unit, School of Life Sciences, The University of Dundee, Dundee DD1 5EH, UK; Faculty of Medicine, Universidad de los Andes, Santiago, Chile; Molecular Biology and Bioinformatics Lab, Program in Molecular Biology and Bioinformatics, Center for Biomedical Research and Innovation (CIIB), Universidad de los Andes, Santiago, Chile

## Abstract

Ubiquitylation regulates all aspects of development, and components are frequently mutated in developmental disorders. Tonne-Kalscheuer Syndrome (TOKAS) is a X-linked multiple congenital anomaly disorder caused by mutations in the E3 ubiquitin ligase RNF12/RLIM and characterized by intellectual disability and urogenital abnormalities. However, the molecular underpinnings of TOKAS remain largely unknown. Here, we show that RNF12 catalytic activity relieves gene repression to drive a transcriptional program required for germ cell development and priming of pluripotent cells towards the germline. A major feature of the RNF12-dependent gametogenesis gene program is a transcriptional feed-forward loop featuring the deubiquitylase *Usp26*/USP26. *Usp26*/USP26 induction stabilises RNF12 to amplify transcriptional responses, which is disrupted by RNF12 TOKAS mutations and USP26 variants identified in patients with fertility defects. In summary, we uncover remarkable synergy within a ubiquitylation cycle that controls expression of key genes required for germ cell development and is disrupted in patients with urogenital abnormalities.

## Introduction

Ubiquitylation is a post-translational modification that serves as a cellular control system by altering the fate and/or function of protein targets (Oh et al., 2018). Depending on ubiquitylation linkage topology, ubiquitylation can target proteins for proteasomal degradation, participate in signalling or promote complex assembly (Kulathu and Komander, 2012). A key role of protein ubiquitylation is regulation of developmental cell fate decisions (Rape, 2018; Werner et al., 2017; Werner and Rape, 2017), and as such ubiquitylation components are frequently mutated in developmental disorders such as intellectual disability (Neri et al., 2018). Tonne-Kalscheuer syndrome (TOKAS) is one such X-linked developmental disorder caused by variants in the E3 ubiquitin ligase RNF12/RLIM (Frints et al., 2019; Hu et al., 2016; Tønne et al., 2015) which impair catalytic activity (Bustos et al., 2018; Frints et al., 2019). TOKAS is characterized by intellectual disability and associated craniofacial abnormalities, as well as syndromic features including urogenital abnormalities, hypogenitalism, velopharyngeal insufficiency and congenital diaphragmatic hernia (Frints et al., 2019; Hu et al., 2016; Tønne et al., 2015). These congenital anomalies can manifest themselves severely in patients, and in the case of diaphragmatic hernia are frequently fatal (Frints et al., 2019).

Recent work by our lab has uncovered an RNF12-dependent signalling pathway that controls expression of neurodevelopmental genes (Bustos et al., 2020), suggesting a potential molecular mechanism by which functional disruption of RNF12 by variants found in TOKAS patients could result in intellectual disability. However, deregulation of neurodevelopmental genes is unlikely to explain other syndromic features characteristic of TOKAS. Therefore, systematic analysis of RNF12-dependent functions in developmental model systems is required to elucidate potential molecular mechanisms that underpin other debilitating syndromic manifestations of TOKAS.

In this paper, we deploy bioinformatics, CRISPR/Cas9 gene editing and biochemical analysis in an embryonic stem cell model to show that RNF12 E3 ubiquitin ligase activity is required for induction of a core gametogenesis gene expression program. We show that gametogenesis gene expression is impaired by an RNF12 TOKAS variant, suggesting that deregulation of this transcriptional program underpins urogenital abnormalities found in TOKAS. RNF12-dependent ubiquitylation and degradation of the REX1 transcriptional repressor unleashes gametogenesis gene expression to drive embryonic stem cell priming towards the germ cell lineage. Furthermore, a major molecular feature of this transcriptional program is induction of a X-linked deubiquitinase, *Usp26*/USP26. RNF12-dependent *Usp26*/USP26 expression drives a biochemical interaction between USP26 and RNF12 resulting in RNF12 stabilisation, thereby establishing a functional feed-forward loop that amplifies RNF12-dependent transcriptional responses. Finally, we show that RNF12 stabilisation and downstream signalling are disrupted by *USP26* gene variants identified from individuals with azoospermia, including sertoli-cell only syndrome. Therefore, our results uncover molecular synergy within the ubiquitylation cycle that controls gametogenesis gene expression and is disrupted in patients with urogenital abnormalities and/or fertility defects.

## Results

### RNF12 E3 ubiquitin ligase activity drives a gametogenesis gene expression program

RNF12 is mutated in the developmental disorder TOKAS, which is characterised by intellectual disability and related craniofacial abnormalities. TOKAS patients suffer other congenital anomalies, including defects in urogenital development/hypogenitalism. In order to investigate molecular mechanisms that may underpin syndromic features of TOKAS, we analysed our previously reported RNA-SEQ dataset (Bustos et al., 2020) to determine developmental gene expression signatures that are positively regulated by RNF12 in male mouse embryonic stem cells (mESCs) cultured in Leukemia Inhibitory Factor (LIF) and Fetal Bovine Serum (FBS; Figure 1A). Amongst RNAs that are induced by WT RNF12 (Figure 1A), Gene Ontology (GO) term analysis identifies significant enrichment of RNAs relating to reproduction, sexual development and gametogenesis (Figure 1B). RNF12-dependent gene expression is also enriched on the X- and Y-chromosomes (Figure 1C), which are known genetic centres of reproductive biology (Burgoyne, 1987; Wang et al., 2001). As RNF12 has not previously been implicated in fertility, gametogenesis and reproduction, and RNF12 TOKAS mutations cause reproductive and urogenital abnormalities, we prioritised this transcriptional program for further investigation.

**Figure 1.**
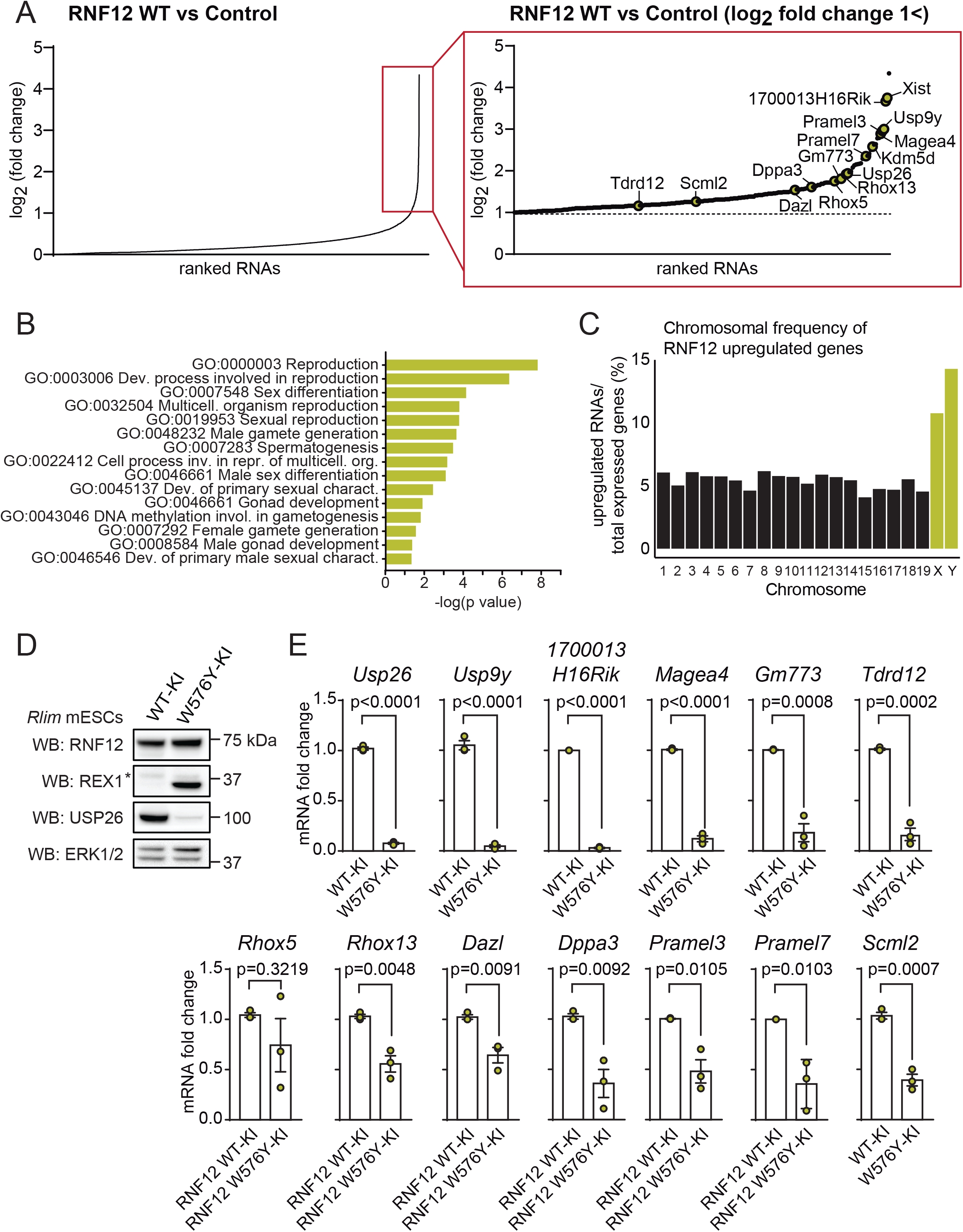
RNF12 E3 ubiquitin ligase activity drives a gametogenesis gene expression program. A) RNA-sequencing (RNA-SEQ) data comparing RNF12-deficient mouse embryonic stem cells (mESCs; *Rlim*^−/y^) transfected with either empty vector (Control) or those expressing wild-type (WT) RNF12 was reported previously (Bustos et al., 2020). RNAs were ranked according to fold-change increase in expression upon RNF12 reconstitution (left). Ranking of RNAs whose expression is increased >2-fold upon RNF12 reconstitution (right). RNAs implicated in reproduction/germ cell development are highlighted. B) Gene Ontology (GO) terms enriched in RNAs whose expression is significantly enriched by RNF12 reconstitution. C) Chromosomal frequency of RNF12 upregulated genes. Percentage of RNAs that are significantly upregulated by RNF12 compared to the total number of expressed genes detected in the RNA-SEQ analysis was calculated for each chromosome. D) Control RNF12 wild-type knock-in (WT-KI) and RNF12 catalytic mutant knock-in (W576Y-KI) mESCs were lysed and RNF12, REX1 and USP26 level determined by immunoblotting. ERK1/2 was used as a loading control. * = non-specific band. E) RNF12 WT-KI and W576Y-KI mESCs were analysed for levels of the indicated mRNAs by qRT-PCR analysis. Data represented as mean ± S.E.M. (n=3). Statistical significance was determined by Student’s T-test; confidence level 95%. *Gapdh* was used as housekeeping control.

We first sought to demonstrate that these RNAs are regulated by endogenous RNF12 E3 ubiquitin ligase activity. A cohort of the most dynamically regulated RNAs implicated in gametogenesis, including *Usp26, Usp9y, Tdrd12, Dazl, Dppa3*, was identified and is summarised (Table 1). RNA expression was analysed in control RNF12 knock-in mESCs (WT-KI) and RNF12 knock-in mESCs harbouring a catalytic mutant that impairs E3 ubiquitin ligase activity (W576Y-KI) (Bustos et al., 2020) leading to accumulation of ubiquitylated RNF12 substrates including the REX1/ZFP42 transcriptional repressor (Figure 1D). Gametogenesis RNAs identified in our RNA-SEQ dataset are expressed in mESCs cultured in LIF/FCS, but this is significantly reduced in RNF12 W576Y-KI mESCs with the exception of *Rhox5* (Figure 1E). Importantly, RNF12 is co-expressed with USP26, DAZL and DPPA3 in mouse testis (Figure S1) suggesting that this relationship extends beyond the stem cell model system. Therefore, our data indicate that RNF12 E3 ubiquitin ligase activity promotes expression of genes involved in the process of gametogenesis, and this might be relevant for urogenital abnormalities observed in TOKAS patients.

**Table 1.**
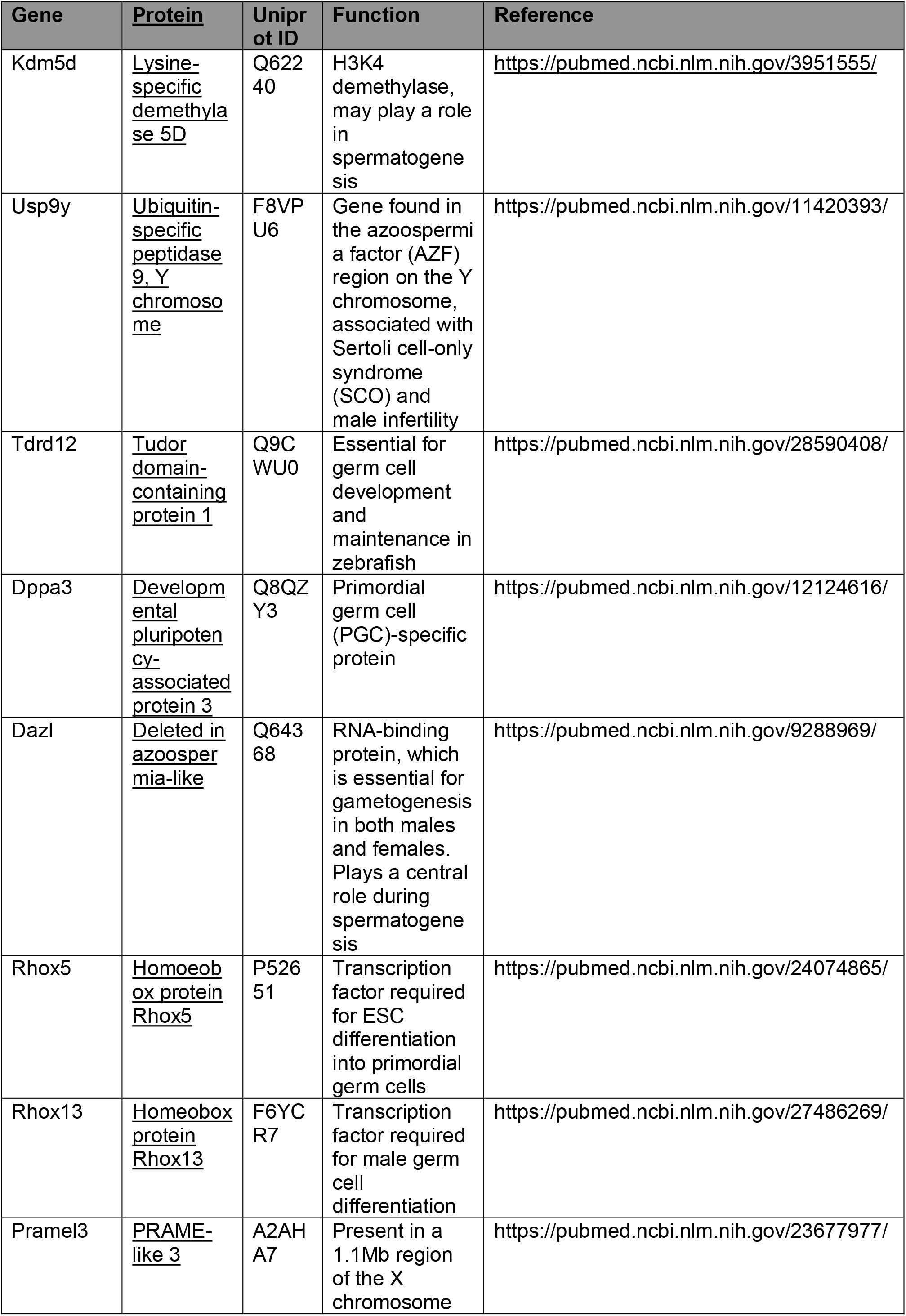
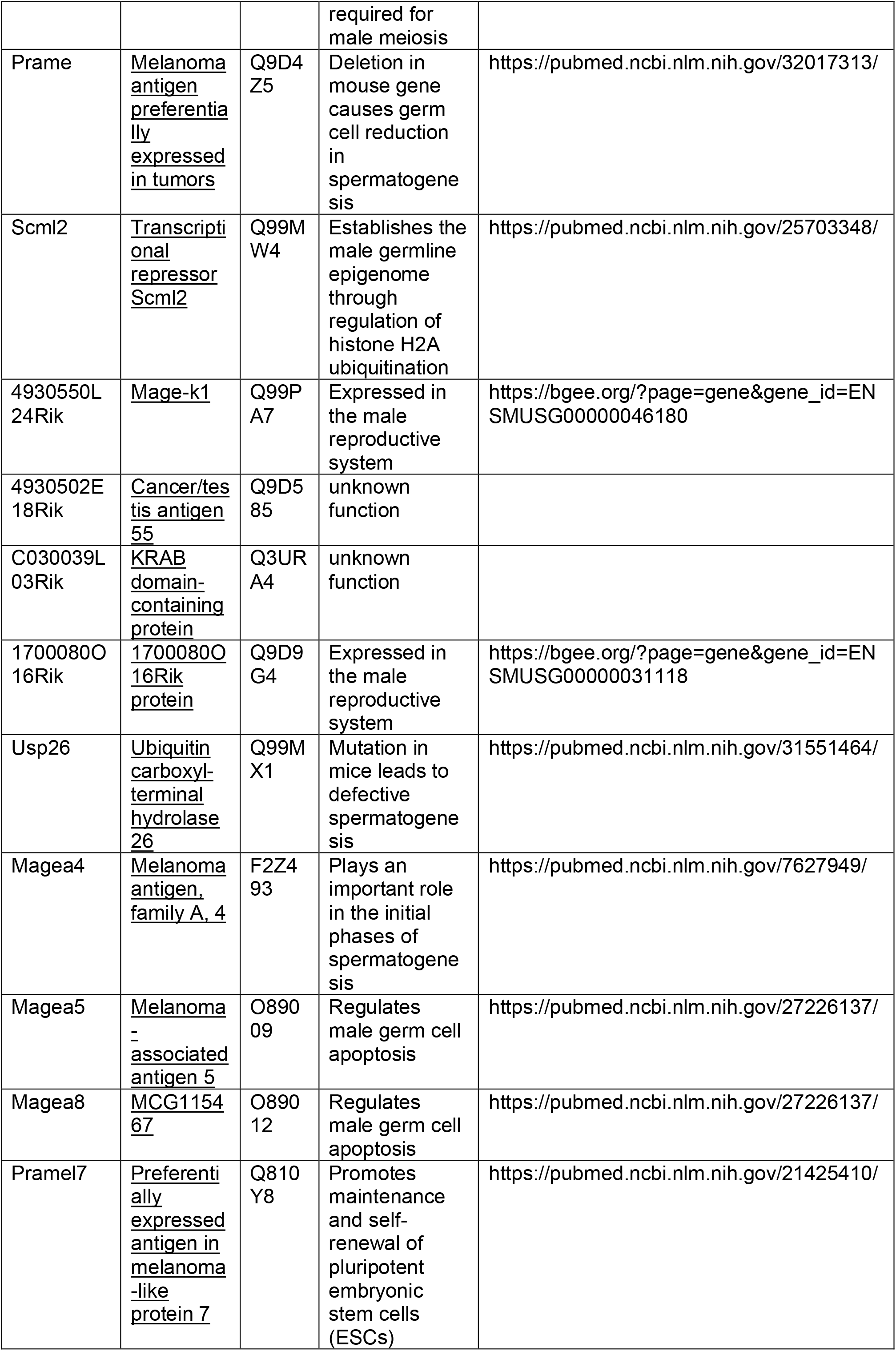
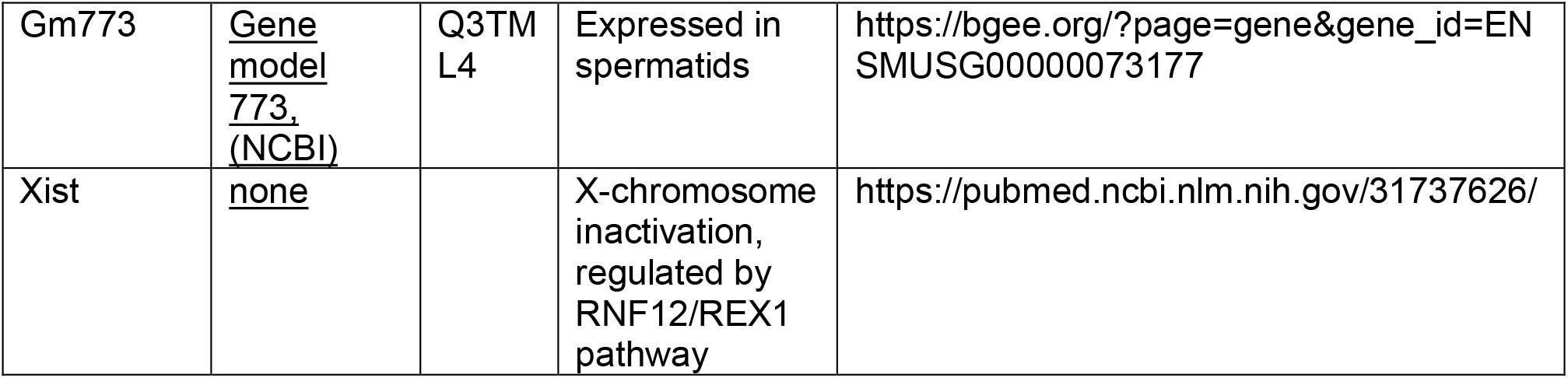
Summary of RNF12-dependent Gametogenesis Genes.

| Gene | Protein | Uniprot ID | Function | Reference |
| --- | --- | --- | --- | --- |
| Kdm5d | <u>Lysine-specific demethylase 5D</u> | Q62240 | H3K4 demethylase, may play a role in spermatogenesis | https://pubmed.ncbi.nlm.nih.gov/3951555/ |
| Usp9y | <u>Ubiquitin-specific peptidase 9, Y chromosome</u> | F8VP U6 | Gene found in the azoospermia factor (AZF) region on the Y chromosome, associated with Sertoli cell-only syndrome (SCO) and male infertility | https://pubmed.ncbi.nlm.nih.gov/11420393/ |
| Tdrd12 | <u>Tudor domain-containing protein 1</u> | Q9C WU0 | Essential for germ cell development and maintenance in zebrafish | https://pubmed.ncbi.nlm.nih.gov/28590408/ |
| Dppa3 | <u>Developmental pluripotency-associated protein 3</u> | Q8QZ Y3 | Primordial germ cell (PGC)-specific protein | https://pubmed.ncbi.nlm.nih.gov/12124616/ |
| Dazl | <u>Deleted in azoospermia-like</u> | Q64368 | RNA-binding protein, which is essential for gametogenesis in both males and females. Plays a central role during spermatogenesis | https://pubmed.ncbi.nlm.nih.gov/9288969/ |
| Rhox5 | <u>Homeobox protein Rhox5</u> | P52651 | Transcription factor required for ESC differentiation into primordial germ cells | https://pubmed.ncbi.nlm.nih.gov/24074865/ |
| Rhox13 | <u>Homeobox protein Rhox13</u> | F6YC R7 | Transcription factor required for male germ cell differentiation | https://pubmed.ncbi.nlm.nih.gov/27486269/ |
| Pramel3 | <u>PRAME-like 3</u> | A2AH A7 | Present in a 1.1Mb region of the X chromosome required for male meiosis | https://pubmed.ncbi.nlm.nih.gov/23677977/ |
| Prame | <u>Melanoma antigen preferentially expressed in tumors</u> | Q9D4Z5 | Deletion in mouse gene causes germ cell reduction in spermatogenesis | https://pubmed.ncbi.nlm.nih.gov/32017313/ |
| Scml2 | <u>Transcriptional repressor Scml2</u> | Q99MW4 | Establishes the male germline epigenome through regulation of histone H2A ubiquitination | https://pubmed.ncbi.nlm.nih.gov/25703348/ |
| 4930550L24Rik | <u>Mage-k1</u> | Q99PA7 | Expressed in the male reproductive system | https://bgee.org/?page=gene&gene_id=ENSMUSG00000046180 |
| 4930502E18Rik | <u>Cancer/testis antigen 55</u> | Q9D585 | unknown function |  |
| C030039L03Rik | <u>KRAB domain-containing protein</u> | Q3UR A4 | unknown function |  |
| 1700080O16Rik | <u>1700080O16Rik protein</u> | Q9D9G4 | Expressed in the male reproductive system | https://bgee.org/?page=gene&gene_id=ENSMUSG00000031118 |
| Usp26 | <u>Ubiquitin carboxyl-terminal hydrolase 26</u> | Q99MX1 | Mutation in mice leads to defective spermatogenesis | https://pubmed.ncbi.nlm.nih.gov/31551464/ |
| Magea4 | <u>Melanoma antigen, family A, 4</u> | F2Z493 | Plays an important role in the initial phases of spermatogenesis | https://pubmed.ncbi.nlm.nih.gov/7627949/ |
| Magea5 | <u>Melanoma-associated antigen 5</u> | O89009 | Regulates male germ cell apoptosis | https://pubmed.ncbi.nlm.nih.gov/27226137/ |
| Magea8 | <u>MCG115467</u> | O89012 | Regulates male germ cell apoptosis | https://pubmed.ncbi.nlm.nih.gov/27226137/ |
| Pramel7 | <u>Preferentially expressed antigen in melanoma-like protein 7</u> | Q810Y8 | Promotes maintenance and self-renewal of pluripotent embryonic stem cells (ESCs) | https://pubmed.ncbi.nlm.nih.gov/21425410/ |
| Gm773 | <u>Gene model 773, (NCBI)</u> | Q3TM L4 | Expressed in spermatids | https://bgee.org/?page=gene&gene_id=ENSMUSG00000073177 |
| Xist | <u>none</u> |  | X-chromosome inactivation, regulated by RNF12/REX1 pathway | https://pubmed.ncbi.nlm.nih.gov/31737626/ |

### RNF12-dependent degradation of the REX1 transcription factor controls gametogenesis gene expression

Next, we set out to identify the mechanism by which RNF12 drives gametogenesis gene expression. The REX1 transcriptional repressor has been identified as a key RNF12 substrate in developmental systems (Bach, 2012; Bustos et al., 2020; Gontan et al., 2012; Gontan et al., 2018). Ubiquitylation and degradation of REX1 relieves transcriptional repression to promote expression of the long non-coding RNA *Xist*, which in turn initiates imprinted X-chromosome inactivation (Gontan et al., 2012; Gontan et al., 2018). We therefore tested whether RNF12 promotes gametogenesis gene expression by relieving REX1-mediated transcriptional repression. To this end, we took advantage of a series of CRISPR/Cas9 mESC lines (Bustos et al., 2020) comprising wild-type (*Rlim*^+/y^), RNF12 gene knockout (*Rlim*^−/y^) and RNF12: REX1 double gene knockout (*Rlim*^−/y^: *Zfp42*^−/−^) (Figure 2A). RNF12 gene knockout leads to increased REX1 levels in these cells, which is reversed by REX1 gene disruption (Figure 2A). Protein expression from two major RNF12-dependent gametogenesis genes identified in our RNA-SEQ analysis, USP26 and DPPA3, is suppressed by RNF12 gene knockout, but restored to levels similar to that of wild-type mESCs by RNF12/REX1 double knockout (Figure 2A). These data suggest that ubiquitylation and degradation of the REX1 transcriptional repressor is a primary mechanism by which RNF12 drives gametogenesis gene expression.

**Figure 2.**
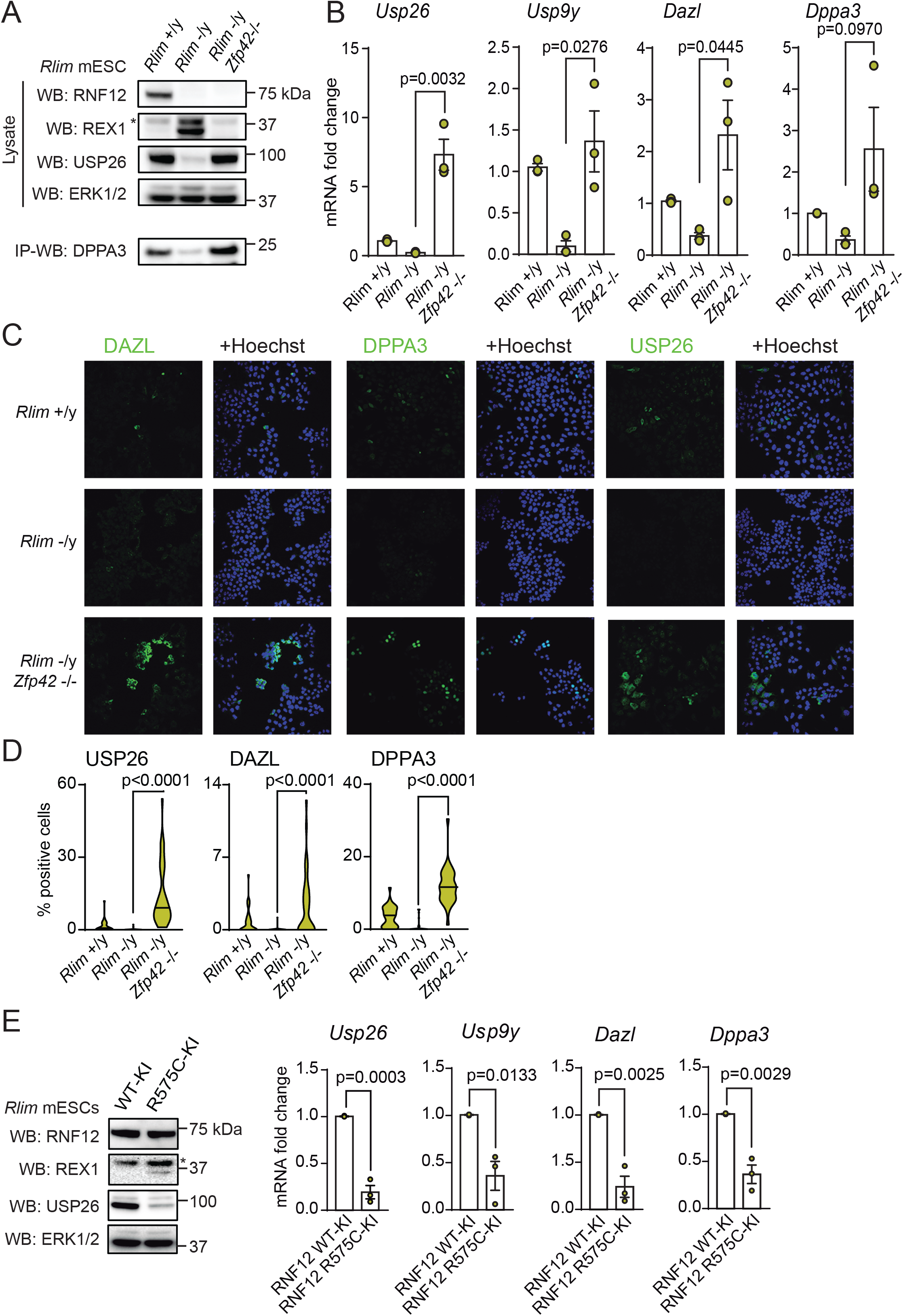
RNF12 induces gametogenesis genes and germ-cell priming via REX1 degradation. A) Control RNF12 wild-type (*Rlim*^−/y^), RNF12-deficient (*Rlim*^−/y^) and RNF12: REX1-deficient (*Rlim*^−/y^: *Zfp42*^−/−^) were lysed and RNF12, REX1, USP26 and DAZL levels determined by immunoblotting. ERK1/2 was used as a loading control. * = non-specific band. B) *Rlim*^−/y^, *Rlim*^−/y^ and *Rlim*^−/y^: *Zfp42*^−/−^ mESCs were analysed for levels of the indicated mRNAs by qRT-PCR analysis. Data represented as mean ± S.E.M. (n=3). Statistical significance was determined by Student’s T-test; confidence level 95%. *Gapdh* was used as housekeeping control. C) *Rlim*^−/y^, *Rlim*^−/y^ and *Rlim*^−/y^: *Zfp42*^−/−^ mESCs were analysed for DAZL, DPPA3 and USP26 expression by immunofluorescence. Hoechst DNA stain is included as a control. D) *Rlim*^−/y^, *Rlim*^−/y^ and *Rlim*^−/y^: *Zfp42*^−/−^ mESCs were analysed for DAZL, DPPA3 and USP26 expression by immunofluorescence. Percentage of DAZL, DPPA3 or USP26 positive cells was determined from >30 fields of view using the number of positive cells divided by the total number of cells per field of view (defined as number of Hoechst positive nuclei). E) Control RNF12 wild-type knock-in (WT-KI) and RNF12 TOKAS mutant knock-in (R575C-KI) mESCs were lysed and RNF12, REX1 and USP26 levels determined by immunoblotting. ERK1/2 was used as a loading control. * = non-specific band (Left). RNF12 WT-KI and R575C-KI mESCs were analysed for levels of the indicated mRNAs by qRT-PCR (Right). Data represented as mean ± S.E.M. (n=3). Statistical significance was determined by Student’s T-test; confidence level 95%. *Gapdh* was used as housekeeping control.

We then determined whether REX1 is the key RNF12 substrate to drive expression of other key gametogenesis mRNAs. As predicted, *Usp26*, *Dazl*, *Usp9y and Dppa3* mRNA levels are suppressed by RNF12 gene knockout (Figure 2B). However, in RNF12/REX1 double knockout mESCs, mRNA expression in each case is restored to levels either similar to or above that of wild-type mESCs (Figure 2B). These results confirm that RNF12-dependent relief of REX1 transcriptional repression plays a wider role in regulation of a gametogenesis gene expression program.

### The RNF12-REX1 axis controls embryonic stem cell priming towards the germ cell lineage

Our results thus far indicate that RNF12-mediated REX1 degradation governs expression of germ cell-specific genes. Thus, we explored the impact of the RNF12-REX1 axis on “priming” of mESCs towards the germ cell lineage. Germ cell priming is a relatively rare event in mESCs cultured in LIF/FBS, as few cells stain positive for gametogenesis gene markers DAZL, DPPA3 and USP26, respectively (Figure 2C). This low level of mESC germ-cell priming is further reduced by RNF12 gene knockout (Figure 2C). However, double knockout of RNF12 and REX1 leads to an increase in the number of mESCs staining positive for DAZL, DPPA3 and USP26 gametogenesis markers (Figure 2C), as confirmed by automated quantification of the percentage of positive cells (Figure 2D). These data indicate that RNF12-mediated REX1 degradation plays a crucial role in controlling the ability of mESCs to prime towards the germ cell lineage, and that genetic disruption of REX1 may lead to unrestricted capacity of mESCs to undergo germ cell lineage priming.

### Gametogenesis genes are disrupted by an RNF12 variant identified from Tonne-Kalscheuer Syndrome (TOKAS) patients

As TOKAS is characterised by syndromic anomalies including urogenital defects with hypogenitalism, we sought to determine the impact of RNF12 TOKAS mutation on RNF12-dependent regulation of gametogenesis genes. To this end, we took advantage of mESCs harbouring a knock-in the mouse equivalent of the human R599C mutation identified in TOKAS patients (RNF12 R575C-KI) (Bustos et al., 2018). Compared to control RNF12 knock-in mESCs (RNF12 WT-KI), RNF12 R575C-KI mESCs display impaired RNF12 E3 ubiquitin ligase activity, causing accumulation of REX1 substrate (Figure 2E). Interestingly, RNF12 R575C-KI mESCs also display reduced levels of USP26 protein, suggesting that TOKAS mutation may interfere with gametogenesis gene expression. Indeed, analysis of *Usp26*, *Dazl*, *Usp9y and Dppa3* gametogenesis mRNAs confirms that all are significantly disrupted in RNF12 R575C-KI mESCs compared to control RNF12 WT-KI mESCs (Figure 2E). These data demonstrate that RNF12 TOKAS mutation interferes with a gametogenesis gene expression program, suggesting that disrupted germ-cell specific gene transcription may underpin urogenital abnormalities observed in TOKAS patients.

### USP26 counteracts RNF12 auto-ubiquitylation to suppress proteasomal degradation

We next sought to uncover key molecular features of the RNF12-dependent gametogenesis gene expression program. A major facet of the ubiquitin system is E3 ubiquitin ligase auto-ubiquitylation leading to “self-destruction”, which often has important regulatory consequences (de Bie and Ciechanover, 2011). We therefore explored the possibility that RNF12 is regulated by auto-ubiquitylation. Indeed, WT RNF12 is ubiquitylated, as determined by tandem-ubiquitin binding element (TUBE) enrichment of ubiquitylated proteins followed by immunoblotting (Figure 3A). Although RNF12 ubiquitylation is not impacted by mutation of 4 of 6 lysine residues present in mouse RNF12, ubiquitylation is lost when all potential acceptor lysine residues are mutated to arginine (RNF12 All K-R), or E3 ubiquitin ligase activity is mutationally inactivated (RNF12 W576Y; Figure 3A). Furthermore, RNF12 auto-ubiquitylation includes K48-linked ubiquitin chains, which are primarily associated with targeting for proteasomal degradation (Chau et al., 1989), as determined by probing RNF12 HA immunoprecipitates with a K48 specific antibody (Figure 3B) or using MUD1 ubiquitin binding elements, which are selective for K48-linked chains (Trempe et al., 2005) (Figure 3C). Importantly, RNF12 auto-ubiquitylation leads to self-destruction, as WT RNF12 is degraded much more rapidly than a non-ubiquitylatable RNF12 mutant (RNF12 All K-R) or an RNF12 E3 ubiquitin ligase deficient mutant (RNF12 W576Y; Figure 3D). These data therefore suggest that RNF12 auto-ubiquitylation modulates cellular levels of RNF12.

**Figure 3.**
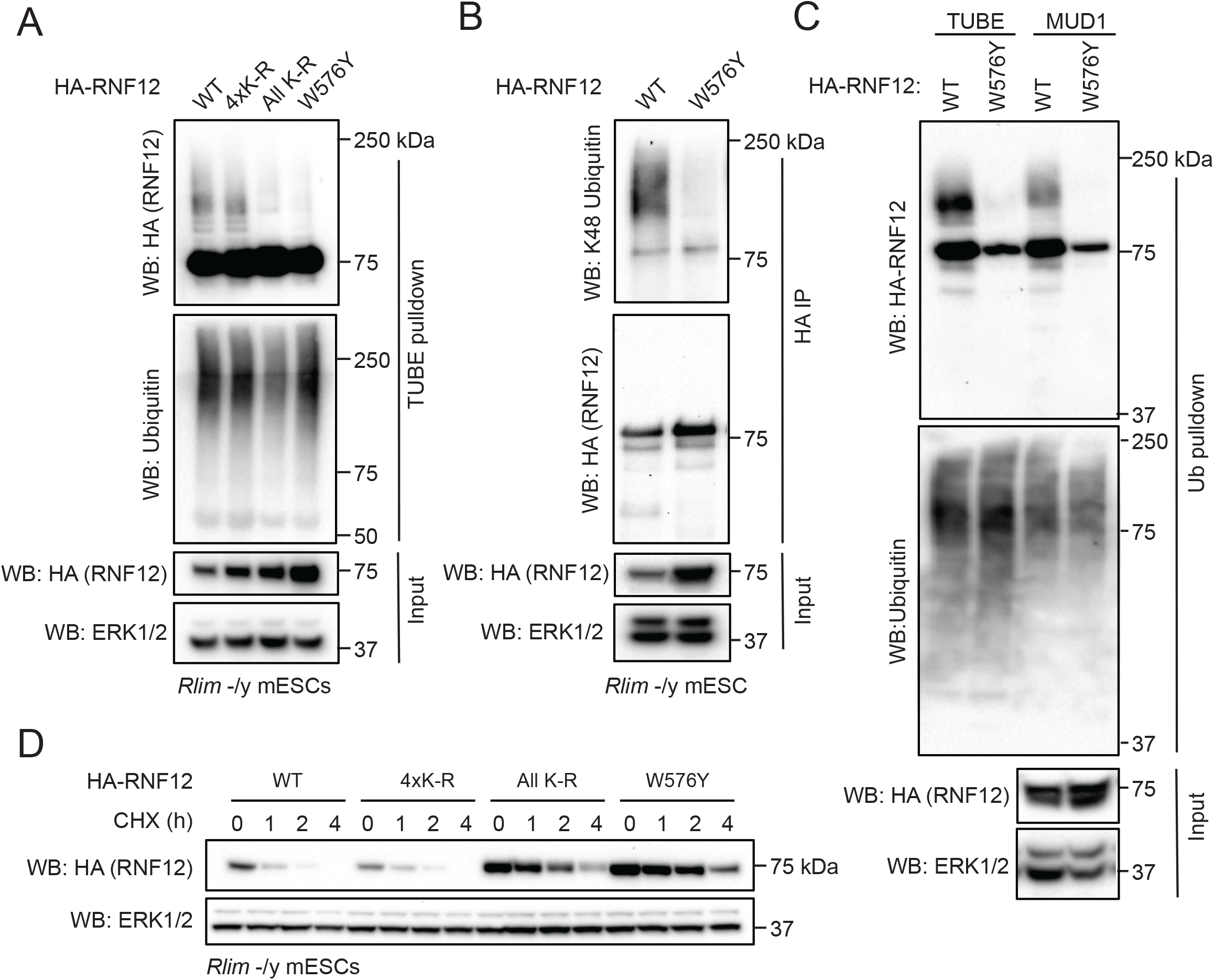
RNF12 auto-ubiquitylation drives rapid proteasomal degradation. A) *Rlim*^−/y^ mESCs were transfected with HA-tagged RNF12 WT, RNF12 K71/526/544/558R (4xK-R), RNF12 K9/71/526/544/558/561R (All K-R) or RNF12 catalytic mutant (W576Y) and ubiquitylated proteins isolated by TUBE pulldown. HA-RNF12 and ubiquitin levels in TUBE pull-downs and HA-RNF12 and ERK1/2 levels in input were determined by immunoblotting. B) *Rlim*^−/y^ mESCs were transfected with HA-tagged RNF12 WT or RNF12 W576Y and RNF12 isolated by HA immunoprecipitation (HA-IP). K48-linked ubiquitin and HA-RNF12 levels in HA-IP and HA-RNF12 and ERK1/2 levels in input were determined by immunoblotting. C) *Rlim*^−/y^ mESCs were transfected with HA-tagged RNF12 WT or RNF12 W576Y and all ubiquitylated proteins isolated by TUBE pulldown or K48-linked ubiquitylated proteins isolated by MUD1 ubiquitin binding domain pull-down (HA-IP). HA-RNF12 and ubiquitin levels in ubiquitin pull-downs and HA-RNF12 and ERK1/2 levels in input were determined by immunoblotting. D) *Rlim*^−/y^ mESCs were transfected with HA-tagged RNF12 WT, K71/526/544/558R (4xK-R), RNF12 K9/71/526/544/558/561R (All K-R) or RNF12 catalytic mutant (W576Y) and protein stability determined by cycloheximide (CHX) chase for the indicated times. HA-RNF12 and ERK1/2 levels were determined by immunoblotting.

As shown previously, the *Usp26*/USP26 deubiquitylase is a major transcriptional target of the RNF12-REX1 signalling pathway, which could in principle modulate ubiquitin signalling via elevated USP26 levels and increased protein de-ubiquitylation. This prompted us to explore the possibility that USP26 induction acts to stabilise RNF12 by preventing “self-destruction”. We took advantage of RNF12 knockout mESCs (*Rlim*^−/y^), which express no endogenous RNF12 and low levels of USP26, to reconstitute RNF12 signalling. As shown previously, RNF12 has a relatively short half-life due to auto-ubiquitylation and proteasomal degradation (Figure 4A). Expression of human USP26 (hUSP26) stabilises RNF12 over a 4 h time course (Figure 4A). This is dependent on proteasomal degradation, as RNF12 stabilisation by USP26 is not further augmented by proteasomal inhibition in the presence of MG132 (Figure 4B). Importantly, hUSP26 is expressed at similar levels to endogenous mUSP26 in these experiments (Figure S2A, compare USP26 staining for endogenous and expressed mUSP26, and FLAG staining for mUSP26 and hUSP26). RNF12 stabilisation is also highly specific to USP26 and its close relative USP29 (Figure 4C), as expression of the more distantly related USP38 or broad specificity deubiquitylating enzyme USP2 do not stabilise RNF12 in a comparable manner (Figure 4C). Like hUSP26, hUSP29 and mUSP38 are localised to the nucleus, at least in part (Figure S2B). Taken together, these data indicate that USP26 and the related USP29 specifically drive RNF12 stabilisation, which may have an impact on RNF12 function.

**Figure 4.**
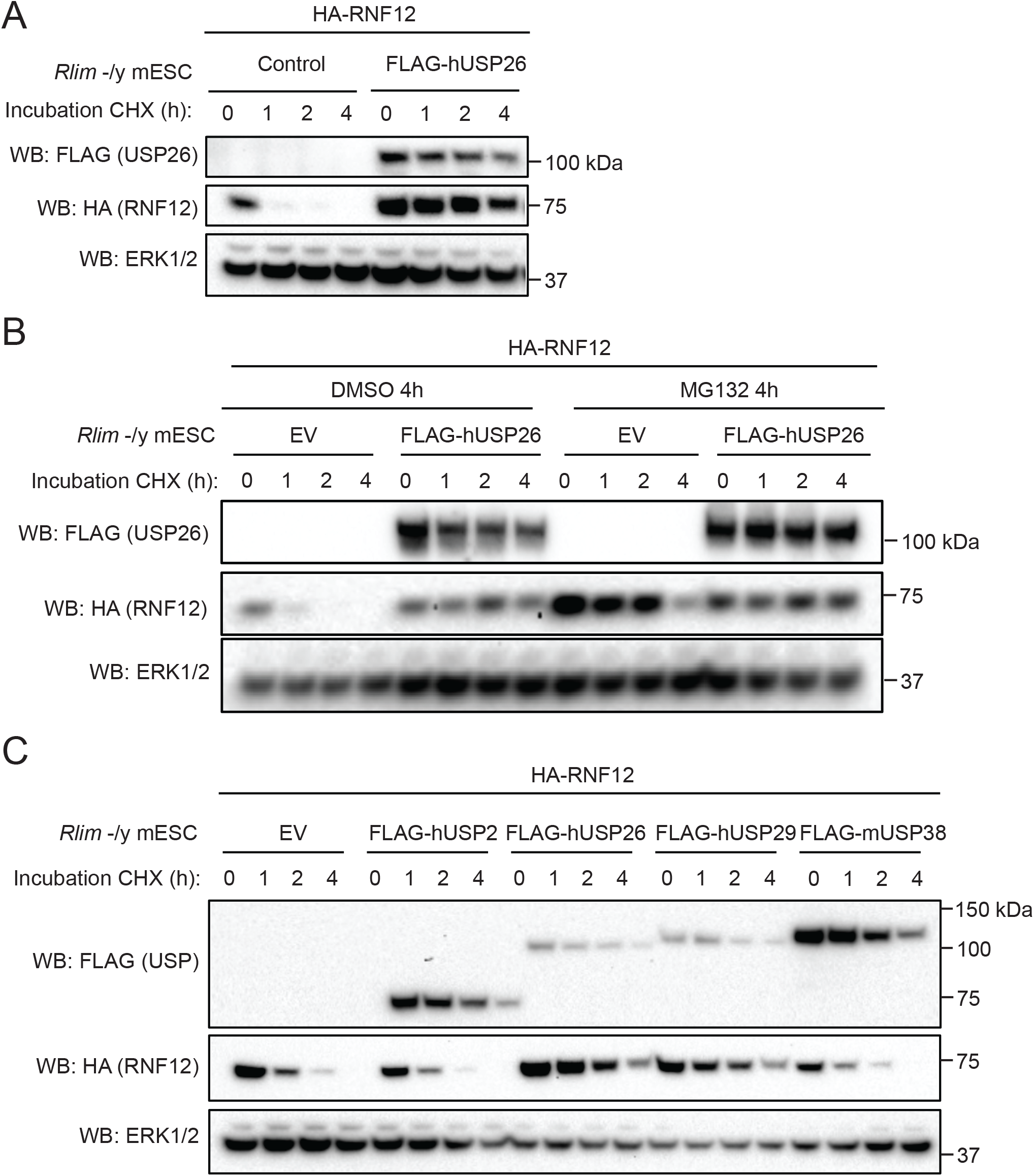
USP26 induction protects RNF12 from rapid proteasomal degradation. A) *Rlim*^−/y^ mESCs were transfected with empty vector or FLAG-tagged human USP26 and HA-tagged RNF12 WT and protein stability determined by cycloheximide (CHX) chase for the indicated times. FLAG-USP26, HA-RNF12 and ERK1/2 levels were determined by immunoblotting. B) *Rlim*^−/y^ mESCs were transfected with empty vector or FLAG-tagged human USP26 and HA-tagged RNF12 WT, treated for DMSO or 10 μ M MG132 for 4h and protein stability determined by cycloheximide (CHX) chase for the indicated times. FLAG-USP26, HA-RNF12 and ERK1/2 levels were determined by immunoblotting. C) *Rlim*^−/y^ mESCs were transfected with empty vector or FLAG-tagged human USP2, USP26, USP29 or USP38 and protein stability determined by cycloheximide (CHX) chase for the indicated times. FLAG-USP26, HA-RNF12 and ERK1/2 levels were determined by immunoblotting.

### USP26 may counteract RNF12 auto-ubiquitylation by preventing RNF12 self-association

These findings prompted us to explore the mechanism by which USP26 regulates RNF12. Firstly, we addressed whether mUSP26 and RNF12 form a biochemical complex in mESCs. Gel filtration analysis indicates that RNF12 and mUSP26 co-elute at a molecular weight 200-500kDa (Figure 5A), suggesting that RNF12 and mUSP26 may form a complex. Co-immunopreciptation analysis reveals that FLAG-hUSP26 specifically interacts with HA-RNF12 (Figure 5B). We also show that hUSP26 is predominantly co-localised to the nucleus with RNF12 (Figure 5C), confirming that RNF12 and USP26 interact in mESCs.

**Figure 5.**
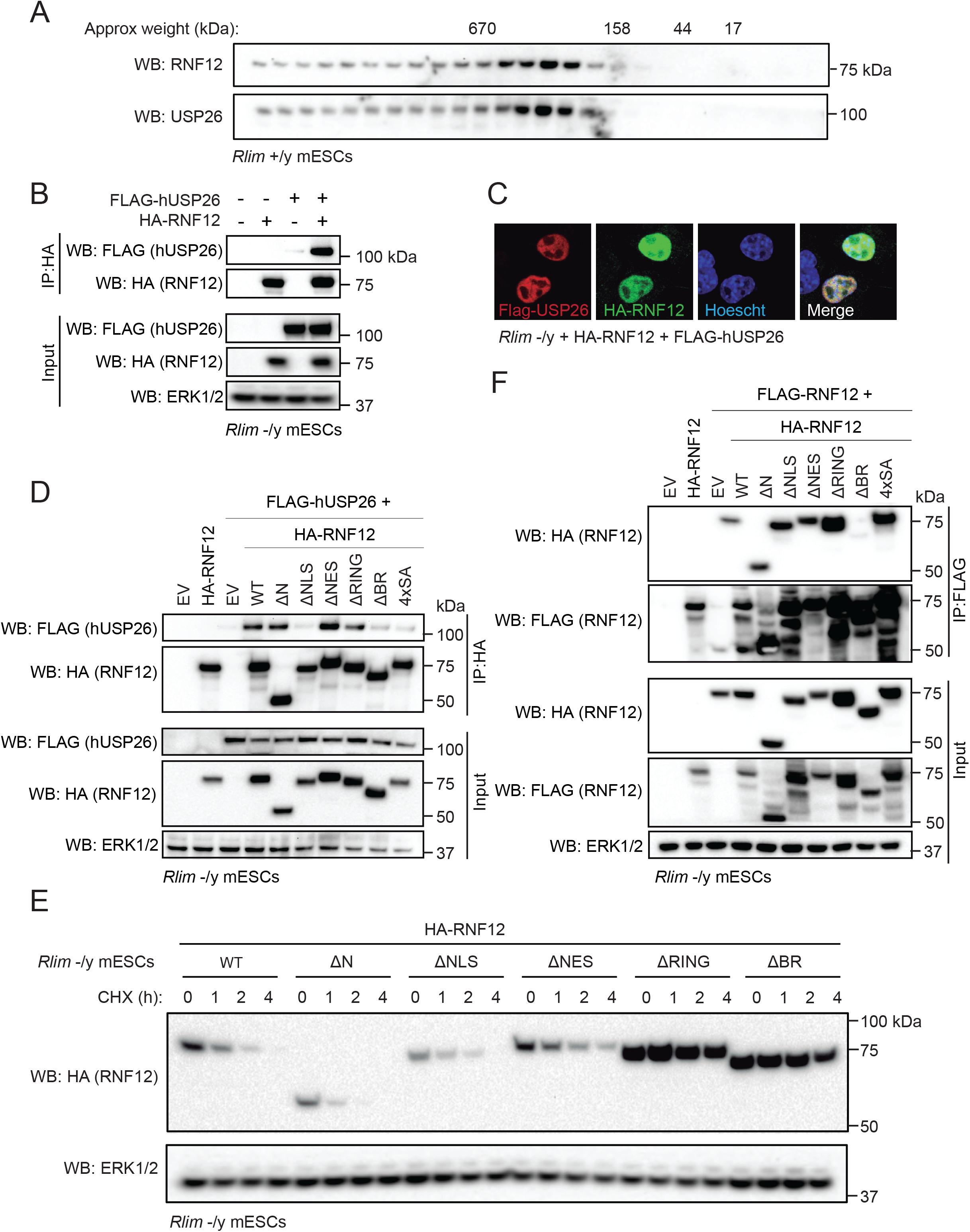
USP26 complexes with RNF12 via the basic region that mediates RNF12 self-association. A) RNF12 WT (*Rlim*^+/y^) mESCs were lysed and extract was analysed by Superose 6 column size-exclusion chromatography. USP26 and RNF12 levels in each fraction from fraction 5 onwards determined by immunoblotting. B) *Rlim*^−/y^ mESCs were transfected with empty vector, FLAG-tagged human USP26 and/or HA-tagged RNF12 WT and RNF12 isolated by HA immunoprecipitation. FLAG-USP26, HA-RNF12 and ERK1/2 levels were determined by immunoblotting. C) *Rlim*^−/y^ mESCs were transfected with FLAG-tagged human USP26 and HA-tagged RNF12 WT and RNF12 and USP26 localisation determined by immunofluorescence. Hoechst DNA stain is shown as a control for nuclei staining. D) *Rlim*^−/y^ mESCs were transfected with empty vector or FLAG-tagged human USP26 and HA-tagged RNF12 WT (1-600), RNF12 Δ1-206 (ΔN), RNF12 Δ206-226 (ΔNLS), RNF12 Δ502-513 (ΔNES) RNF12 Δ546-587 (ΔRING), RNF12 Δ326-423 (ΔBR) or RNF12 S212/214/227/229A (4xSA) and RNF12 isolated by HA immunoprecipitation. FLAG-USP26, HA-RNF12 and ERK1/2 levels were determined by immunoblotting. E) *Rlim*^−/y^ mESCs were transfected with HA-tagged RNF12 WT (1-600), RNF12 Δ1-206 (ΔN), RNF12 Δ206-226 (ΔNLS), RNF12 Δ502-513 (ΔNES) RNF12 Δ546-587 (ΔRING), RNF12 Δ326-423 (ΔBR) and protein stability determined by cycloheximide (CHX) chase for the indicated times. HA-RNF12 and ERK1/2 levels were determined by immunoblotting. F) *Rlim*^−/y^ mESCs were transfected with empty vector or both FLAG and HA-tagged RNF12 WT (1-600), RNF12 Δ1-206 (ΔN), RNF12 Δ206-226 (ΔNLS), RNF12 Δ502-513 (ΔNES) RNF12 Δ546-587 (ΔRING), RNF12 Δ326-423 (ΔBR) or RNF12 S212/214/227/229A (4xSA) and RNF12 isolated by HA immunoprecipitation. FLAG-USP26, HA-RNF12 and ERK1/2 levels were determined by immunoblotting.

We then sought to define RNF12 motif(s) that are responsible for USP26 interaction. Deletion analysis of key RNF12 functional regions (Figure S3A) indicates that the N-terminus predicted leucine-zipper like region (LZL), the catalytic RING domain (RING) and the nuclear export signal (NES) are dispensable for hUSP26 interaction (Figure 5D). Deletion of the RNF12 nuclear localisation signal (NLS) or mutation of the NLS phosphorylation sites (4xSA) disrupts interaction with hUSP26 (Figure 5D), presumably because these mutants are mis-localised to the cytosol when ectopically expressed (Figure S3B) (Bustos et al., 2020; Jiao et al., 2013). However, the RNF12 basic region (BR) deletion mutant, which is co-localised to the nucleus with hUSP26 (Figure S3B), fails to interact with hUSP26. We therefore conclude that the RNF12 BR mediates a biochemical interaction with hUSP26.

Our results suggest that the RNF12 BR plays a key role in RNF12 regulation by recruiting USP26. We hypothesised that deletion of the RNF12 BR will destabilise RNF12 by disrupting the interaction with USP26, thereby preventing RNF12 deubiquitylation. To test this prediction, we examined the impact of RNF12 BR deletion on stability. RNF12 WT and LZL and NLS or NES deletion mutants are relatively unstable due to auto-ubiquitylation (Figure 5E). As expected, deletion of the RNF12 RING domain leads to RNF12 stabilisation (Figure 5E), presumably because this truncated protein lacks catalytic activity. Deletion of the RNF12 BR also stabilises RNF12, and to a similar extent as RING deletion (Figure 5E, quantified in Figure S3C), despite the failure of this construct to interact with USP26 (Figure 5C). This suggests that the RNF12 BR is somehow involved in RNF12 autoubiquitylation, which is consistent with previous data that the RNF12 BR is involved in catalytic activity via an undefined mechanism (Bustos et al., 2018).

RING-type E3 ubiquitin ligases frequently function as dimers (Metzger et al., 2014), suggesting that RNF12 auto-ubiquitylation may require RNF12 self-association. As deletion of the RNF12 BR stabilises the protein, we tested whether the BR provides a platform for RNF12 self-association and auto-ubiquitylation. Indeed, deletion of the RNF12 basic region prevents RNF12 self-association (Figure 5F), suggesting that RNF12 self-assembly may be a key mechanism for auto-ubiquitylation. Furthermore, our demonstration that the RNF12 BR mediates interaction with USP26 suggests that USP26 engagement by RNF12 might indirectly prevent RNF12 self-destruction by occluding a critical site for self-association and auto-ubiquitylation.

### *USP26* variants from azoospermia patients disrupt RNF12 stabilisation and gametogenesis gene expression

USP26 is specifically expressed in testes (Figure S1) (Lin et al., 2011; Wang et al., 2001; Zhang et al., 2009) and is a key component within the RNF12-dependent germ cell specific transcriptional program that is disrupted in by *RNF12/RLIM* TOKAS variants. Interestingly, *USP26* variants are implicated in azoospermia, including sertoli-cell only syndrome (A Paduch et al., 2005; Arafat et al., 2020; Stouffs et al., 2005), which prompts the hypothesis that *USP26* variants may lead to deregulation of RNF12-dependent functions. *USP26* variants are largely found clustered around a nuclear localisation sequence (NLS) and within the USP catalytic domain (Figure 6A), with at least one implicated in disruption of catalytic activity (Liu et al., 2018). Thus, we prioritised a panel of *USP26* variants including frequently reported mutations (Figure 6A) to investigate their impact on USP26 function in RNF12 regulation. In each case, we find that *USP26* variants disrupt expression (Figure 6B), whilst all *USP26* variants tested, including those surrounding the NLS, localise correctly to the nucleus (Figure 6C). These data suggest that disrupted protein expression may be a pathogenic mechanism in fertility patients harbouring USP26 variants.

**Figure 6.**
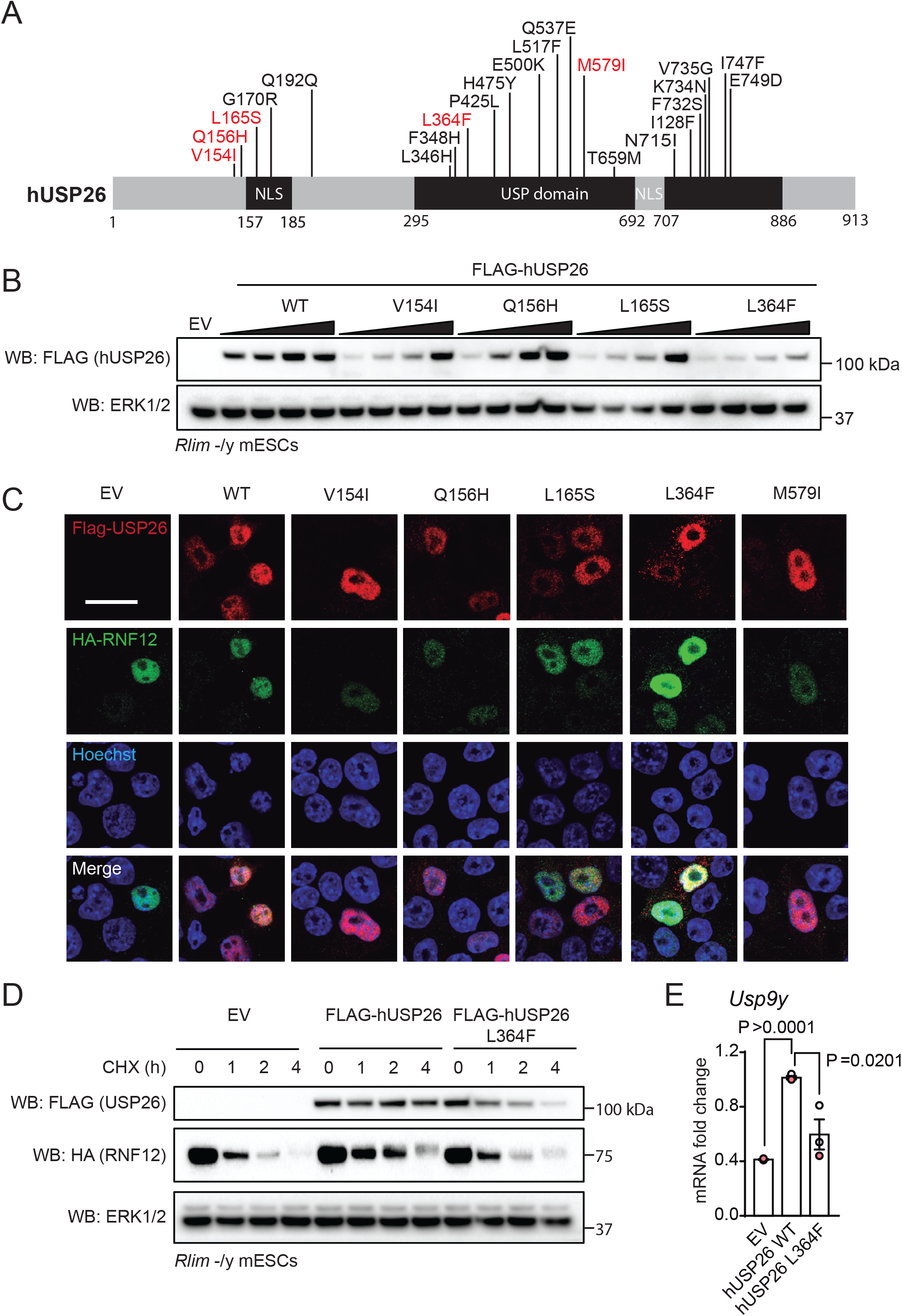
USP26 variants identified in azoospermia patients disrupt RNF12 stabilisation and downstream functions. A) Schematic diagram of human USP26 showing positions of functional regions and variants associated with azoospermia. Variants used in subsequent studies are highlighted in red. B) *Rlim*^−/y^ mESCs were transfected empty vector or increasing amounts of FLAG-tagged USP26 WT or the indicated azoospermia variants. FLAG-USP26 and ERK1/2 levels were determined by immunoblotting. C) *Rlim*^−/y^ mESCs were transfected with FLAG-tagged human USP26 WT or indicated azoospermia variant and HA-tagged RNF12 WT and HA-RNF12 and FLAG-USP26 localisation determined by immunofluorescence. Hoechst DNA stain is shown as a control for nuclei staining. D) *Rlim*^−/y^ mESCs were transfected with HA-tagged RNF12 and either empty vector or FLAG-tagged human USP26 WT or L364F and protein stability determined by cycloheximide (CHX) chase for the indicated times. FLAG-USP26, HA-RNF12 and ERK1/2 levels were determined by immunoblotting. E) *Rlim*^−/y^ mESCs were transfected with HA-tagged RNF12 and either empty vector or FLAG-tagged human USP26 WT or L364F and levels of *Usp9y* mRNA determined by qRT-PCR analysis. Data represented as mean ± S.E.M. (n=3). Statistical significance was determined by Student’s T-test; confidence level 95%. *Gapdh* was used as housekeeping control.

Our finding that *USP26* variants found in fertility patients disrupt protein expression prompts the hypothesis that these variants may impact on RNF12 deubiquitylation and stabilisation. We addressed this possibility using USP26 L364F, a variant within the USP catalytic domain (Figure 6A) which exhibits severely impaired expression (Figure 6B), as an exemplar. As shown previously, RNF12 is rapidly degraded via auto-ubiquitylation, and is stabilised by expression of WT hUSP26 (Figure 6D). However, hUSP26 L364F is also impaired in its ability to promote RNF12 stabilisation (Figure 6D), indicating that USP26 azoospermia mutants not only disrupt expression but may interfere with regulation of RNF12 stability. Of note, hUSP26 WT is a relatively long-lived protein (T1/2 >4h), but the USP26 L364F variant is rapidly degraded (Figure 6D), consistent with reduced overall protein levels of USP26 azoospermia variants.

Finally, we sought to address the impact of the hUSP26 L364F azoospermia-associated variant on RNF12 downstream functions. To this end, we employed the *Usp9y* as a sensitive assay for RNF12-dependent regulation of gametogenesis gene expression. Whilst WT USP26 results in a significant increase in *Usp9y* expression, this effect is significantly impaired by expression of the azoospermia-associated USP26 L364F variant (Figure 6E). Taken together, our data indicate that USP26 amplifies RNF12 signalling to promote gametogenesis gene expression, and this is functionally disrupted by *USP26* variants found in fertility patients (Figure 7).

**Figure 7.**
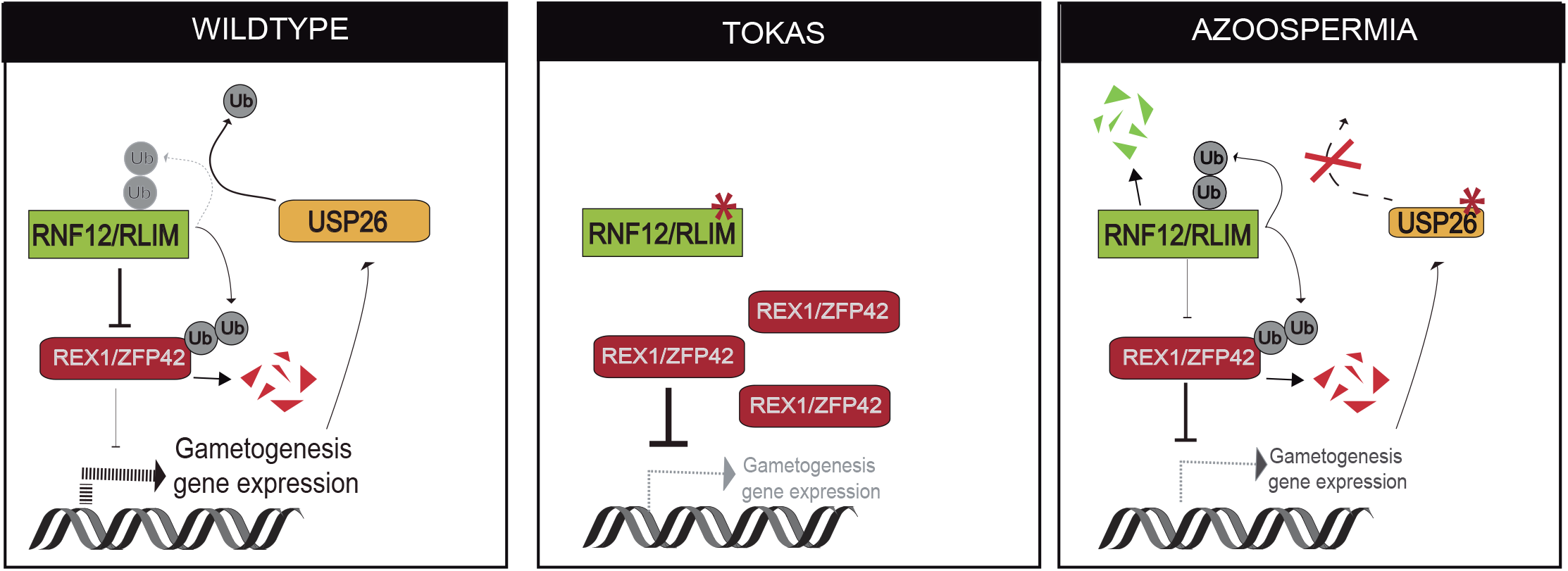
The RNF12-USP26 signalling axis controls gametogenesis gene expression and is disrupted in developmental disorders with urogenital dysfunction.

## Discussion

Ubiquitylation is critical for regulating many human developmental processes and as such ubiquitylation components are mutationally disrupted in a number of devastating human developmental disorders. However, the molecular underpinnings by which ubiquitylation functions in these developmental and disease systems remain almost completely unknown. Here, we uncover a ubiquitylation signalling network involving the RING-type E3 ubiquitin ligase RNF12 and a deubiquitylating enzyme, USP26, which drives expression of gametogenesis genes. We show that RNF12 E3 ubiquitin ligase activity drives *Usp26* gene expression, leading elevated USP26 levels and RNF12 stabilisation. This creates a feed-forward amplification loop to drive RNF12-dependent signalling and a gametogenesis gene expression program. Our results provide detailed molecular insight into the complex interplay within the ubiquitin system in this essential developmental process, and how this is deregulated in urogenital disorders including TOKAS and azoospermia (Figure 7). Of particular interest is the fact that an E3 ubiquitin ligase employs transcriptional induction of a deubiquitylating enzyme to amplify its own function, which is an unprecedented molecular mechanism for activation of a ubiquitin signalling pathway. However, this fits well with previous understanding of how cell-fate decisions are executed and reinforced during development, which frequently involves amplification and negative feedback loops within the signalling and transcriptional machinery to confer robust cellular decision-making (Freeman, 2000).

Our data shows that RNF12-mediated REX1 degradation controls expression of genes involved in germ cell priming, which complements the recent finding that RNF12/RLIM is required for spermiogenesis in mice (Wang et al., 2020a). We build on this important observation by defining the molecular mechanisms by which RNF12 regulates gametogenesis genes and find that USP26 and the closely related USP29 may also be involved in fertility by regulating RNF12. Although USP26 is mutated in patients with fertility defects, there is conflicting data regarding the role of USP26 in gametogenesis and fertility (Felipe-Medina et al., 2019; Sakai et al., 2019; Tian et al., 2019; Wang et al., 2020b) and USP29 is not required for male fertility in mice (Huang et al., 2019). However, our data show that USP26 and USP29 could in principle function redundantly to regulate RNF12 stability, and so a compound USP26/29 knock-out approach may be required to address this. Interestingly, *Usp26* but not *Usp29* gene expression is regulated by RNF12, suggesting that USP29 may play a constitutive role in this system.

Finally, the mechanism by which USP26 stabilises RNF12 remains to be determined. USP26 may be recruited to RNF12 to directly catalyse deubiquitylation and stabilisation. However, the fact that USP26 engages RNF12 within a region that also promotes RNF12 self-association, leading to turnover, suggests that USP26 might function via a more complex mechanism whereby it disrupts RNF12 self-association to actively prevent auto-ubiquitylation. This will be resolved by development of active recombinant USP26 for use in *in vitro* RNF12 auto-ubiquitylation and substrate ubiquitylation assays.

## Experimental Procedures

### Cell culture

Male mouse embryonic stem cells (mESCs, CCE line) were cultured in ES-DMEM media (DMEM base, 5% knockout serum replacement (KO serum) (v/v), 2 mM glutamine, 0.1 mM minimum essential media (MEM) nonessential amino acids, 1 mM sodium pyruvate, 50 U penicillin, 50 μg/ml streptomycin, 10% FBS (v/v), 0.1 mM β-mercaptoethanol; supplemented with 20 ng/mL LIF (Medical Research Council Protein Phosphorylation and Ubiquitin Unit Reagents and Services (MRC-PPU R&S)) on plates coated with 0.1% gelatin at 5% CO_2_ and 37°C.

### Transfection and plasmids

mESCs were transfected using Lipofectamine LTX (Thermo Fisher Scientific) according to the manufacturer’s instructions. pCAGGS puro plasmids (Table S1) were generated by the MRC-PPU R&S and verified by DNA sequencing (MRC-PPU DNA Sequencing & Services) using Applied Biosystems Big-Dye Ver 3.1 chemistry on an Applied Biosystems model 3730 automated capillary DNA sequencer. All cDNA clones can be found at the MRC-PPU R&S website http://mrcppureagents.dundee.ac.uk/.

### Pharmacological inhibition

Small molecule inhibitors/compounds used are listed in Table S2. MG132 and cycloheximide treatments were at a final concentration of 10 μM and 350 μM, respectively.

### CRISPR Cas9 genome editing

Genome editing was conducted using CRISPR (Clustered Regularly Interspaced Short Palindromic Repeat)/Cas9 system (Doudna and Charpentier, 2014). *Rlim*^−/y^ RNF12 knockout, RNF12 wild-type knock-in (WT-KI) and R575C-KI (Bustos et al., 2018), RNF12 W576Y-KI and *Rlim*^−/y^:*Zfp42*^−/−^ RNF12:REX1 double knock-out (Bustos et al., 2020) mESCs were described previously using guide RNA sequences summarised in Table S3. For knock-in experiments a third vector containing the donor DNA sequence followed by an IRES-EGFP cassette was co-expressed (Table S4). CRISPR/Cas9 plasmid DNAs were transfected into mESCs using Lipotectamine (LTX) according to the manufacturer’s instructions and cultured for 24 h. Transfected mESCs were selected with 3 μg/ml puromycin for 48 h and then subjected into single-cell sorting using a FACS instrument into gelatinized 96-well cell culture plates. Candidate clones were analysed by immunoblotting and genomic DNA sequencing to confirm gene editing. Sequencing was carried out by MRC-PPU DNA Sequencing & Services.

### Immunofluorescence

2×10^5^ cells/cm^2^ were plated into 12-well plates containing a single gelatin-coated coverslip per well. For transfected mESCs, this procedure was conducted 24 h post-transfection. Cells were left to attach for 24 h, media removed, and cells washed twice with PBS. Cells were then fixed to coverslips using 4% (w/v) paraformaldehyde (PFA), diluted in PBS and incubated for 20 mins at room temperature in the dark. Fixed cells were then washed three times with PBS and permeabilized in 0.5% Triton X-100 (w/v) diluted in PBS for 5 mins at room temperature. Permeabilised cells were blocked using 1% (w/v) fish gelatin diluted in PBS for at least 30 mins at room temperature inside a humid chamber. Cells were then stained with primary antibody (Table S5) in 1% (w/v) fish gelatin diluted in PBS for 2 h at room temperature and in a humid chamber. After washing with PBS three times, cells were stained with secondary antibody tagged to fluorophore (Table S5) diluted in 1% fish gelatin for 1 h at room temperature in a humid chamber and in darkness. DNA was stained using

0.1 μg/ml Hoechst diluted in PBS for 5 mins at room temperature in the dark. Coverslips were mounted on microscope slides using microscopy grade mounting media and left to dry at room temperature in the dark for 24 h before imaging. Images were acquired in a Zeiss 710 confocal microscope using Zen software (Zeiss).

### Protein extraction

mESCs were harvested using lysis buffer (20 mM Tris-HCl (pH 7.4), 150 mM NaCl, 1 mM EDTA, 1% (v/v) NP-40, 0.5% (w/v) sodium deoxycholate, 10 mM β-glycerophosphate, 10 mM sodium pyrophosphate, 1 mM NaF, 2 mM Na_3_VO_4_, and 0.1 U/ml Complete Protease Inhibitor Cocktail Tablets (Roche)). Mouse organs were harvested from 19-week-old C57BL/6J mice and snap frozen in liquid nitrogen, resuspended in lysis buffer and lysed using a Polytrone PT 1200 E homogeniser (Kinematica AG). Mouse studies were approved by the University of Dundee ethical review committee, and further subjected to approved study plans by the Named Veterinary Surgeon and Compliance Officer (Dr. Ngaire Dennison) and performed under a UK Home Office project licence in accordance with the Animal Scientific Procedures Act (ASPA, 1986). Protein concentration of protein extracts was determined using BCA Protein Assay Kit (Pierce) according to manufacturer’s directions. Protein concentration was calculated using a standard BSA protein curve.

### Immunoprecipitation

For RNF12 and DPPA3 immunoprecipitations, 20 μl protein G agarose beads were washed three times in lysis buffer and incubated with 1 mg of mESC lysate and 2 μg of RNF12 or DPPA3 antibody (see Table S6) overnight at 4 °C. For FLAG- or HA-tag pulldowns, 10 μl of pre-coupled FLAG-M2 agarose or 20 μl HA-Sepharose beads were washed three times with lysis buffer and incubated with 1 mg of mESC lysate overnight at 4°C. Modified haloalkane dehalogenase (HALO)-tagged tandem ubiquitin binding element (HALO-TUBE) beads and HALO-MUD1 beads were produced as described (Emmerich and Cohen, 2015). 1 mg of mESC lysate was mixed with 40 μl of HALO-TUBE or HALO-MUD1 beads and incubated overnight at 4 °C on a rotating wheel. In all cases, beads were washed three times with lysis buffer containing 500 mM NaCl. At each step, beads were centrifuged at 2000 rpm for 2 mins and supernatant was discarded. Finally, proteins bound to the beads were eluted by the addition of LDS sample buffer and boiling the mixture 5 mins at 95 °C.

### Size Exclusion Chromatography

Size exclusion chromatography (SEC) running buffer (50 mM Tris-HCl (pH 7.5), 10% Glycerol (v/v), 150 mM NaCl, 1 mM DTT was freshly prepared and degassed by passing through a 0.45 μm PVDF filter (Millipore). 24 h prior to use, an ÄKTA™ pure protein purification instrument was equilibrated using SEC running buffer. A single confluent 15 cm plate of mESCs was harvested in SEC cell collection buffer (1 mM EGTA, 1 mM EDTA in PBS). Detached cells were collected into a Falcon tube and centrifuged 1000 rpm for 5 minutes at 4°C. Supernatant was discarded and pellet resuspended in 5 volumes of lysis buffer (20 mM Tris-HCl (pH 7.4), 150 mM NaCl, 1 mM EDTA, 1% NP-40 (v/v), 0.5% sodium deoxycholate (w/v), 10 mM β-glycerophosphate, 10 mM sodium pyrophosphate, 1 mM NaF, 2 mM Na_3_VO_4_ and 0.1 U/ml Complete Protease Inhibitor Cocktail Tablets (Roche)) and incubated for 15 mins on ice. After lysis, mixture was subjected to centrifugation at 14,000 rpm for 15 mins at 4 °C. Supernatant was then collected and passed through a 0.45 μm filter. Finally, clarified sample was mixed with Gel Filtration Standards (molecular weight range 670 – 1.35 kDa #1511901, Bio-Rad) and injected into an ÄKTA™ pure protein purification instrument loaded with a Superose 6 column through which a standard size exclusion chromatography protocol was run and 30 fractions collected per sample.

### Western blotting

NuPAGE ^TM^ 4-12% SDS-PAGE gels were transferred onto PVDF membrane and incubated with primary antibody (see Table S6) diluted in 5% (w/v) non-fat milk buffer in TBS-T overnight at 4 °C. After incubation, membranes were washed 3 times with TBS-T buffer (20 mM Tris-HCl (pH 7.5), 150 mM NaCl supplemented with 0.2% (v/v) Tween-20 (Sigma Aldrich)) and incubated for 1 h with secondary horseradish Peroxidase (HRP)-conjugated antibodies (see Table S6) at room temperature. Finally, membranes were washed 3 times with TBS-T and subjected to chemiluminescence detection with Immobilon Western Chemiluminescent HRP (Horseradish Peroxidase) Substrate (Merck Millipore) using a Gel-Doc XR+ System (BioRad). Acquired images were analysed using ImageLab software (BioRad).

### RNA extraction and qRT-PCR

mESCs were seeded in 6-well plates and left to grow 24-48 h until confluent. RNA was extracted using E.Z.N.A.® MicroElute Total RNA Kit (Omega Bio-Tek) according to manufacturer’s instructions. RNA samples were subjected to reverse transcription using iScript™ cDNA Synthesis Kit (BioRad) according to the manufacturer’s instructions. qPCR primer sequences were identified from the PrimerBank database (https://pga.mgh.harvard.edu/primerbank) or designed using Primer3 software with a melting temperature between 58-62 °C. For each primer, the integrity and specificity were confirmed in silico by NCBI Primer-Blast software (https://www.ncbi.nlm.nih.gov/tools/primer-blast). All primers are 20-24 bases long with an overlap of seven bases at the intron/exon boundary producing an amplicon of 100-300 bases. Primers were supplied by Invitrogen or Thermo Scientific according to availability. qPCR primer sequences are listed in Table S7. qPCR reactions were carried out using SsoFast EvaGreen Supermix (BioRad) or TB Green Premix Ex Taq (Takara). In either case, each sample consisted of a 10 μl reaction containing 1 μl cDNA with 400 nM forward and reverse primers, 5 μl SYBR Green and nuclease free water. Each sample was prepared in technical duplicate. qPCR and absorbance detection were carried out in a CFX384 real-time PCR system (BioRad). The ΔΔCt method, also known as the Pfaffl method (Pfaffl, 2001) was used to calculate relative RNA levels with *Gapdh* expression as a loading control. Data were analysed in Excel software and plotted in GraphPad Prism v.8.00 software.

### RNA-sequencing (RNA-SEQ) and gene ontology (GO-term) analysis

RNA-SEQ data for *Rlim*^−/y^ mESCs expressing either empty vector or RNF12 WT was described previously (Bustos et al., 2020) (see Gene Expression Omnibus (GEO) accession GSE149554). Reads were aligned using Spliced Transcripts Alignment to a Reference (STAR) software. Differential gene expression was estimated using DESeq2 package and further statistical analysis and plot generation were performed with The SARTools R package. Gene Ontology (GO) analysis were carried out using the GO stat R package.

### Data analysis

Data is presented as mean ± S.E.M. with individual points representing a single biological replicate. In qPCR experiments two technical replicates were run per sample and averaged. Immunofluorescence images were processed using ImageJ (ImageJ) and Photoshop CS5.1 (Adobe) software. The percentage of cells expressing a certain protein was calculated as the ratio between cells positive for protein expression (containing antibody fluorescence) and the total number of cells (containing DNA fluorescence provided by Hoechst stain). Parameters were quantified using Fiji (ImageJ) software. Graphs were created using Prism software (GraphPad). In all cases, statistical significance was determined through ANOVA followed by Tukey’s post hoc test or student’s T-test using Prism software (GraphPad) and significant differences were considered when p<0.05.

## Acknowledgements

We thank Prof Wolf Reik and Dr Julia Spindel (Babraham Institute) and Prof Vicky Cowling (University of Dundee) for helpful discussions, and Prof Vicky Cowling and Dr Joana Silva (University of Dundee) for mouse tissue extracts. G.M.F and F.B are supported by a Wellcome Trust/Royal Society Sir Henry Dale Fellowship (211209/Z/18/Z) and a Medical Research Council New Investigator Award (MR/N000609/1). A.S-F. is supported by a MRC-PPU prize studentship. G.N. is supported by research grant ANID/FONDECYT 11190998

## Author Contributions

A.S-F, F.B. and G.M.F. conceived the study and designed the experiments. F.B. & A.S-F. performed experiments. R.T. cloned plasmid DNA. G.N. analysed data and prepared figures. A.S-F., F.B. and G.M.F. wrote the paper.

## References

A Paduch, D., Mielnik, A., and Schlegel, P.N. (2005). Novel mutations in testis-specific ubiquitin protease 26 gene may cause male infertility and hypogonadism. In Reproductive BioMedicine Online, pp. 747–754.

Arafat, M., Zeadna, A., Levitas, E., Har Vardi, I., Samueli, B., Shaco-Levy, R., Dabsan, S., Lunenfeld, E., Huleihel, M., and Parvari, R. (2020). Novel mutation in USP26 associated with azoospermia in a Sertoli cell-only syndrome patient. In Molecular Genetics {&} Genomic Medicine (John Wiley {&} Sons, Ltd), pp. e1258.

Bach, I. (2012). Releasing the break on X chromosome inactivation: Rnf12/RLIM targets REX1 for degradation. In Cell research (Nature Publishing Group), pp. 1524–1526.

Burgoyne, P.S. (1987). The role of the mammalian Y chromosome in spermatogenesis. Development 101 Suppl, 133–141.

Bustos, F., Segarra-Fas, A., Chaugule, V.K., Brandenburg, L., Branigan, E., Toth, R., Macartney, T., Knebel, A., Hay, R.T., Walden, H., et al. (2018). RNF12 X-Linked Intellectual Disability Mutations Disrupt E3 Ligase Activity and Neural Differentiation. Cell Rep 23, 1599–1611.

Bustos, F., Segarra-Fas, A., Nardocci, G., Cassidy, A., Antico, O., Davidson, L., Brandenburg, L., Macartney, T.J., Toth, R., Hastie, C.J., et al. (2020). Functional Diversification of SRSF Protein Kinase to Control Ubiquitin-Dependent Neurodevelopmental Signaling. Dev Cell.

Chau, V., Tobias, J.W., Bachmair, A., Marriott, D., Ecker, D.J., Gonda, D.K., and Varshavsky, A. (1989). A multiubiquitin chain is confined to specific lysine in a targeted short-lived protein. Science 243, 1576–1583.

de Bie, P., and Ciechanover, A. (2011). Ubiquitination of E3 ligases: self-regulation of the ubiquitin system via proteolytic and non-proteolytic mechanisms. In Cell death and differentiation (Nature Publishing Group), pp. 1393–1402.

Doudna, J.A., and Charpentier, E. (2014). The new frontier of genome engineering with CRISPR-Cas9. In Science, pp. 1258096.

Emmerich, C.H., and Cohen, P. (2015). Optimising methods for the preservation, capture and identification of ubiquitin chains and ubiquitylated proteins by immunoblotting. In Biochemical and Biophysical Research Communications, pp. 1–14.

Felipe-Medina, N., Gómez-H, L., Condezo, Y.B., Sanchez-Mart\’in, M., Barbero, J.L., Ramos, I., Llano, E., and Pendás, A.M. (2019). Ubiquitin-specific protease 26 (USP26) is not essential for mouse gametogenesis and fertility. In Chromosoma, pp. 237–247.

Freeman, M. (2000). Feedback control of intercellular signalling in development. Nature 408, 313–319.

Frints, S.G.M., Ozanturk, A., Rodr\’iguez Criado, G., Grasshoff, U., de Hoon, B., Field, M., Manouvrier-Hanu, S., E. Hickey, S., Kammoun, M., Gripp, K.W., et al. (2019). Pathogenic variants in E3 ubiquitin ligase RLIM/RNF12 lead to a syndromic X-linked intellectual disability and behavior disorder. In Molecular Psychiatry, pp. 1748–1768.

Gontan, C., Achame, E.M., Demmers, J., Barakat, T.S., Rentmeester, E., van IJcken, W., Grootegoed, J.A., and Gribnau, J. (2012). RNF12 initiates X-chromosome inactivation by targeting REX1 for degradation. In Nature, pp. 386–390.

Gontan, C., Mira-Bontenbal, H., Magaraki, A., Dupont, C., Barakat, T.S., Rentmeester, E., Demmers, J., and Gribnau, J. (2018). REX1 is the critical target of RNF12 in imprinted X chromosome inactivation in mice. In Nature communications (Nature Publishing Group), pp. 4752.

Hu, H., Haas, S.A., Chelly, J., Van Esch, H., Raynaud, M., de Brouwer, A.P.M., Weinert, S., Froyen, G., Frints, S.G.M., Laumonnier, F., et al. (2016). X-exome sequencing of 405 unresolved families identifies seven novel intellectual disability genes. In Molecular psychiatry (Nature Publishing Group), pp. 133–148.

Huang, Z., Khan, M., Xu, J., Khan, T., Ma, H., Khan, R., Hussain, H.M.J., Jiang, X., and Shi, Q. (2019). The deubiquitinating gene Usp29 is dispensable for fertility in male mice. In Science China Life Sciences, pp. 544–552.

Jiao, B., Taniguchi-Ishigaki, N., Güngör, C., Peters, M.A., Chen, Y.-W., Riethdorf, S., Drung, A., Ahronian, L.G., Shin, J., Pagnis, R., et al. (2013). Functional activity of RLIM/Rnf12 is regulated by phosphorylation-dependent nucleocytoplasmic shuttling. In Molecular biology of the cell, pp. 3085–3096.

Kulathu, Y., and Komander, D. (2012). Atypical ubiquitylation - the unexplored world of polyubiquitin beyond Lys48 and Lys63 linkages. Nat Rev Mol Cell Biol 13, 508–523.

Lin, Y.-W., Hsu, T.-H., and Yen, P.H. (2011). Localization of ubiquitin specific protease 26 at blood–testis barrier and near Sertoli cell–germ cell interface in mouse testes. In International Journal of Andrology (John Wiley {&} Sons, Ltd), pp. e368--e377.

Liu, Y.-L., Zheng, J., Mi, Y.-J., Zhao, J., and Tian, Q.-B. (2018). The impacts of nineteen mutations on the enzymatic activity of USP26. In Gene, pp. 292–296.

Metzger, M.B., Pruneda, J.N., Klevit, R.E., and Weissman, A.M. (2014). RING-type E3 ligases: master manipulators of E2 ubiquitin-conjugating enzymes and ubiquitination. Biochim Biophys Acta 1843, 47–60.

Neri, G., Schwartz, C.E., Lubs, H.A., and Stevenson, R.E. (2018). X-linked intellectual disability update 2017. In American journal of medical genetics Part A, pp. 1375–1388.

Oh, E., Akopian, D., and Rape, M. (2018). Principles of Ubiquitin-Dependent Signaling. Annu Rev Cell Dev Biol 34, 137–162.

Pfaffl, M.W. (2001). A new mathematical model for relative quantification in real-time RT-PCR. In Nucleic acids research (Oxford University Press), pp. e45--e45.

Rape, M. (2018). Ubiquitylation at the crossroads of development and disease. Nat Rev Mol Cell Biol 19, 59–70.

Sakai, K., Ito, C., Wakabayashi, M., Kanzaki, S., Ito, T., Takada, S., Toshimori, K., Sekita, Y., and Kimura, T. (2019). Usp26 mutation in mice leads to defective spermatogenesis depending on genetic background. In Scientific Reports, pp. 13757.

Stouffs, K., Lissens, W., Tournaye, H., Van Steirteghem, A., and Liebaers, I. (2005). Possible role of USP26 in patients with severely impaired spermatogenesis. In European Journal of Human Genetics, pp. 336–340.

Tian, H., Huo, Y., Zhang, J., Ding, S., Wang, Z., Li, H., Wang, L., Lu, M., Liu, S., Qiu, S., et al. (2019). Disruption of ubiquitin specific protease 26 gene causes male subfertility associated with spermatogenesis defects in mice†. In Biology of Reproduction, pp. 1118–1128.

Tønne, E., Holdhus, R., Stansberg, C., Stray-Pedersen, A., Petersen, K., Brunner, H.G., Gilissen, C., Hoischen, A., Prescott, T., Steen, V.M., et al. (2015). Syndromic X-linked intellectual disability segregating with a missense variant in RLIM. In European journal of human genetics : EJHG (Nature Publishing Group), pp. 1652–1656.

Trempe, J.-F., Brown, N.R., Lowe, E.D., Gordon, C., Campbell, I.D., Noble, M.E.M., and Endicott, J.A. (2005). Mechanism of Lys48-linked polyubiquitin chain recognition by the Mud1 UBA domain. In The EMBO journal, pp. 3178–3189.

Wang, F., Gervasi, M.G., Bošković, A., Sun, F., Rinaldi, V.D., Yu, J., Wallingford, M.C., Tourzani, D.A., Mager, J., Zhu, L.J., et al. (2020a). Deficient spermiogenesis in mice lacking <em>Rlim</em>. bioRxiv, 2020.2008.2031.275248.

Wang, J., Zhao, X., Hong, R., and Wang, J. (2020b). USP26 deubiquitinates androgen receptor (AR) in the maintenance of sperm maturation and spermatogenesis through the androgen receptor signaling pathway. Adv Clin Exp Med.

Wang, P.J., McCarrey, J.R., Yang, F., and Page, D.C. (2001). An abundance of X-linked genes expressed in spermatogonia. In Nature Genetics, pp. 422–426.

Werner, A., Manford, A.G., and Rape, M. (2017). Ubiquitin-Dependent Regulation of Stem Cell Biology. Trends Cell Biol 27, 568–579.

Werner, A., and Rape, M. (2017). Powering stem cell decisions with ubiquitin. Cell Death Differ 24, 1823–1824.

Zhang, J., Tian, H., Huo, Y.-W., Zhou, D.-X., Wang, H.-X., Wang, L.-R., Zhang, Q.-Y., and Qiu, S.-D. (2009). The expression of Usp26 gene in mouse testis and brain. In Asian journal of andrology (Nature Publishing Group), pp. 478–483.

