## Supplemental information for "A RNF12-USP26 amplification loop promotes germ cell specification and is disrupted in urogenital disorders"

**Supplementary Tables**

**Table S1: Plasmids**

| cDNA | Construct | MRC-PPU R&S NUMBER | Cloned by |
|---|---|---|---|
| Empty vector | pCAGGS puro | DU49023 | Rachel Toth |
| HA-RNF12 WT | pCAGGS puro HA mouse RNF12 | DU50854 | Rachel Toth |
| HA-RNF12 ΔN | pCAGGS puro HA mouse RNF12 A206-V600(end) | DU53408 | Rachel Toth |
| HA-RNF12 ΔNLS | pCAGGS puro HA mouse RNF12 ΔA206-R226 | DU53426 | Rachel Toth |
| HA-RNF12 ΔNES | pCAGGS puro HA mouse RNF12 ΔL502-L513 | DU53405 | Rachel Toth |
| HA-RNF12 ΔRING | pCAGGS puro HA mouse RNF12 M1-L543 | DU53419 | Rachel Toth |
| HA-RNF12 ΔBR | pCAGGS puro HA mouse RNF12 ΔY326-A423 | DU53422 | Rachel Toth |
| HA-RNF12 4XSA | pCAGGS puro HA mouse RNF12 S212/214/227/229A | DU58741 | Rachel Toth |
| HA-RNF12 W576Y | pCAGGS puro HA mouse RNF12 W576Y | DU61086 | Jennifer Crooks |
| HA-RNF12 4xK-R | pCAGGS puro HA mouse RNF12 K71R K526R K544R K558R | DU61130 | Jennifer Crooks |
| HA-RNF12 All K-R | pCAGGS puro HA mouse RNF12 K9R K71R K526R K544R K558R 561R | DU61139 | Jennifer Crooks |
| FLAG-RNF12 WT | pCAGGS puro 3XFLAG mouse Rnf12 | DU49070 | Rachel Toth |
| FLAG-RNF12 ΔN | pCAGGS puro 3FLAG mouse RNF12 A206-V600(end) | DU53409 | Rachel Toth |
| FLAG-RNF12 ΔNLS | pCAGGS puro 3XFLAG mouse RNF12 ΔA206-R226 | DU53416 | Rachel Toth |
| FLAG-RNF12 ΔNES | pCAGGS puro 3XFLAG mouse RNF12 ΔL502-L513 | DU53421 | Rachel Toth |
| FLAG-RNF12 ΔRING | pCAGGS puro 3XFLAG mouse RNF12 M1-L543 | DU53417 | Rachel Toth |
| FLAG-RNF12 ΔBR | pCAGGS puro 3XFLAG mouse RNF12 ΔY326-A423 | DU53418 | Rachel Toth |
| FLAG-RNF12 4XSA | pCAGGS puro 3FLAG mouse RNF12 S212/214/227/229A | DU67399 | Rachel Toth |
| FLAG-USP26 WT | pCAGGS puro 3FLAG mouse USP26 | DU53288 | Rachel Toth |

| | | | |
|---|---|---|---|
| FLAG-USP26 V154I | pCAGGS puro 3FLAG human USP26 V154I | DU61166 | Jennifer Crooks |
| FLAG-USP26 Q156H | pCAGGS puro 3FLAG human USP26 Q156H | DU61169 | Jennifer Crooks |
| FLAG-USP26 L165S | pCAGGS puro 3FLAG human USP26 L165S | DU61170 | Jennifer Crooks |
| FLAG-USP26 L364F | pCAGGS puro 3FLAG human USP26 L364F | DU61171 | Jennifer Crooks |
| FLAG-USP26 M579I | pCAGGS puro 3FLAG human USP26 M579I | DU61188 | Jennifer Crooks |
| FLAG-USP2 | pCAGGS puro 3FLAG human USP2 | DU61200 | Jennifer Crooks |
| FLAG-USP26 | pCAGGS puro 3FLAG human USP26 | DU67207 | Rachel Toth |
| FLAG-USP29 | pCAGGS puro 3FLAG human USP29 | DU61201 | Rachel Toth |
| FLAG-USP38 | pCAGGS puro 3XFLAG mouse USP38 | DU49091 | Rachel Toth |

**Table S2: Chemicals**

| COMPOUND | STOCK CONC (mM) | DILUTION | SOURCE | REF Nº | TARGET |
|---|---|---|---|---|---|
| MG132 | 10 | DMSO | SIGMA | M8699 | Proteasome |
| Cycloheximide | 350 | Ethanol | SIGMA | C4859 | 60S ribosomal unit |

**Table S3: CRISPR/Cas9 guide RNA details**

| mESC CRISPR LINE | TARGET GENE | MRC-PPU R&S PROJECT N° | CONSTRUCT | MRC-PPU R&S DU NUMBER | SENSE /ANTI-SENSE | CRISPR CAS9 TARGETING SEQUENCE |
|---|---|---|---|---|---|---|
| *Rlim*$^{-/y}$ | RNF12/RLIM | CR124 | pBabeD P U6 RLIM (mouse) ex 5 KO Sense A | DU52037 | Sense | GATAAATGTTAA CCGTAACAA |
| | | | pX335 RLIM (mouse) ex5 KO AntiSense A+Cas9n | DU52046 | Anti-sense | GAATCTGAAAT CACCGCTGTT |
| *Rlim*$^{-/y}$: *Zfp42*$^{-/-}$ | REX1/RNF12 | CR670 | pBabeD P U6 ZFP42 (mouse) ex 4 KO Sense A | DU60065 | Sense | GAGGAAGATGG CTTCCCTGA |
| | | | pX335 ZFP42 (mouse) ex4 KO AntiSense A+Cas9n | DU6007 | Anti-sense | GAATCTCACTTT CATCCCGGA |
| RNF12-KI | RNF12/RLIM | CR697 | pBabeD P U6 RLIM (mouse) Cter KI Sense A | DU57881 | Sense | GCAGGGCAGTC TTATCTTCT |
| | | | pX335 RLIM (mouse) Cter KI AntiSense A+Cas9n | DU57891 | Anti-sense | GTGGAATTCTC AGACAACCAG |

**Table S4: Donor constructs for CRISPR/Cas9 knock-in**

| mESC CRISPR KI | MRC-PPU R&S PROJECT Nº | DONOR CONSTRUCT | DU NUMBER |
|---|---|---|---|
| RNF12 WT-KI | CR697 | pMA RLIMm Cter wt control IRES-GFP donor | DU57967 |
| RNF12 W576Y KI | CR697 | pMA RLIMm Cter W576Y IRES-GFP donor | DU60290 |
| RNF12 R575C KI | CR644 | pMA RLIMm Cter R575C IRES-GFP donor | DU57916 |

**Table S5: Antibodies for immunofluorescence**

| ANTIBODY | SOURCE | ORGANISM | REFERENCE | DILUTION |
|---|---|---|---|---|
| DAZL | MRCPPU R&S | SHEEP | S836B | 1:200 |
| DPPA3 | ABCAM | RABBIT | ab19878 | 1:200 |
| USP26 | MRCPPU R&S | SHEEP | SA085 | 1:1000 |
| FLAG | SIGMA | MOUSE | F1804-50 | 1:500 |
| HA | ABCAM | RABBIT | ab9110 | 1:1000 |
| anti-MOUSE AlexaFluor 555nm | LifeTech | DONKEY | A31570 | 1:500 |
| anti-RABBIT AlexaFluor 488nm | LifeTech | GOAT | A11008 | 1:500 |
| anti-RABBIT AlexaFluor 546nm | LifeTech | GOAT | A11010 | 1:500 |
| anti-SHEEP AlexaFluor 488nm | LifeTech | DONKEY | A11015 | 1:500 |

**Table S6: Antibodies for immunoprecipitation and immunoblotting**

| ANTIBODY | SOURCE | ORGANISM | REFERENCE | DILUTION |
|---|---|---|---|---|
| RNF12 | MRCPPU R&S | SHEEP | S691D | 1:1000 |
| REX1 | ABCAM | RABBIT | ab28141 | 1:1000 |
| FLAG-HRP | SIGMA | MOUSE | A8592-2 | 1:1000 |
| HA-HRP | ROCHE | RABBIT | 12013819001 | 1:1000 |
| ERK1/2 | SANTA CRUZ | RABBIT | SC-93 | 1:1000 |
| ERK1/2 | BD | MOUSE | 610408 | 1:1000 |
| UBIQUITIN | DAKO | RABBIT | Z0458 | 1:1000 |
| K48 UBIQUITIN | CST | RABBIT | 4289 | 1:1000 |
| USP26 | MRCPPU R&S | SHEEP | SA085 | 1:1000 |
| DAZL | MRCPPU R&S | SHEEP | S836B | 1:1000 |
| DPPA3 | ABCAM | RABBIT | ab19878 | 1:1000 |
| αSHEEP-HRP | MRCPPU R&S | | | 1:10000 |
| αRABBIT-HRP | CST | GOAT | 7074S | 1:10000 |
| αMOUSE-HRP | CST | HORSE | 7076S | 1:10000 |

**Table S7: qRT-PCR primers**

| PRIMERS | NCBI GENE ID | DIRECTION | SEQUENCE |
|---|---|---|---|
| *170013H16RIK* | 75514 | Forward | GGAGTTGACATTAACCGTGCT |
| | | Reverse | CATTAAGCTGTGCCATTGCATC |
| *Dazl* | 13164 | Forward | TGGACCGAAGCATACAGACAGTGGT |
| | | Reverse | TGATCAGATTTAAGCACTGCCCGAC |
| *Dppa3* | 73708 | Forward | GACCCAATGAAGGACCCTGAA |
| | | Reverse | GCTTGACACCGGGGTTTAG |
| *Gapdh* | 14433 | Forward | CTCGTCCCGTAGACAAAA |
| | | Reverse | TGAATTTGCCGTGAGTGG |
| *Gm773* | 331416 | Forward | TCTGTTTCAGCAGTGGGATTTTGA |
| | | Reverse | AGTGCTTTCAGGCTGTGGACCT |
| *Magea4* | 17140 | Forward | GGCTCACCTATGATGGGATGCT |
| | | Reverse | CTCACTGACACAGTTTCCTTGCG |
| *Pramel3* | 83565 | Forward | CCTTTTGCCTGTCTCCACATTGG |
| | | Reverse | CAGCCAGCATCCTGCCTTAAATC |
| *Pramel7* | 347712 | Forward | GTGAGGAATGAAGTATTGACCGT |
| | | Reverse | TCAGCCATGTGTCTACTCCATC |
| *Rhox5* | 18617 | Forward | ACTCGGAAGAACAGCATGATG |
| | | Reverse | CCCTGGTGCCACTATCCTT |
| *Rhox13* | 73614 | Forward | ACCGCCATTCCACTTCGCAC |
| | | Reverse | ATTGGGCACAGAGGTTGC |
| *Scml2* | 107815 | Forward | ATCTTCCCAGTTGGATGGTG |
| | | Reverse | CTGGGGCCTCTTCTTCATTT |
| *Tdrd12* | 71981 | Forward | GGGCTCTGATTAAGTCCATCATC |
| | | Reverse | ACTTGGCAAAATCGACCAGGA |
| *Usp9y* | 107868 | Forward | ATGGCAGGTTGCACATTCAC |
| | | Reverse | CAGTCCATCTTGATCATTTGG |
| *Usp26* | 83563 | Forward | GCACTGGATGCTAAATGCAA |
| | | Reverse | TGTGCTGAGTGCCTGTCCTA |

#### Supplementary Figure Legends

**Figure S1 (Related to Figure 1).**

**RNF12 E3 ubiquitin ligase activity drives a gametogenesis gene expression programme**.

Indicated mouse tissue extracts were analysed for RNF12, USP26, DAZL, DPPA3 and Actin expression by immunoblotting.

**Figure S2 (Related to Figure 4).**

**Expression and localisation of USP26 and related deubiquitylases**

*A) Rlim*$^{+/y}$ mESCs were transfected with HA-tagged RNF12 and empty vector, and *Rlim*$^{-/y}$ mESCs were transfected with HA-tagged RNF12 and either empty vector, FLAG-tagged human USP26 or FLAG-tagged mouse USP26 WT. Transfected hUSP26/mUSP26 was detected by FLAG or total USP26 immunofluorescence, endogenous mUSP26 was detected by total USP26 immunofluorescence. Hoechst DNA stain is included as a control. B) *Rlim*$^{-/y}$ mESCs were transfected with HA-tagged RNF12 and either empty vector, FLAG-tagged human USP2, FLAG-tagged human USP26, FLAG-tagged human USP29 or FLAG-tagged mouse USP38. Transfected FLAG-USPs were detected by FLAG immunofluorescence, transfected HA-RNF12 was detected by HA immunofluorescence. Hoechst DNA stain is included as a control.

**Figure S3 (Related to Figure 5)**

**USP26 complexes with RNF12 via the basic region that mediates RNF12 self-association.**

A) Schematic of RNF12 deletion and point mutants used in USP26 interaction studies. B) *Rlim*$^{-/y}$ mESCs were transfected with the HA-tagged RNF12 WT (1-600), RNF12 Δ1-206 (ΔN), RNF12 Δ206-226 (ΔNLS), RNF12 Δ502-513 (ΔNES) RNF12 Δ546-587 (ΔRING), RNF12 Δ326-423 (ΔBR). HA-RNF12 localisation was detected by HA immunofluorescence. Hoechst DNA stain is included as a control for nuclei staining. C) *Rlim*$^{-/y}$ mESCs were transfected with HA-tagged RNF12 WT (1-600), RNF12 Δ546-587 (ΔRING) and RNF12 Δ326-423 (ΔBR) and protein levels determined by cycloheximide (CHX) chase for the indicated times. HA-RNF12 and ERK1/2 levels were determined by immunoblotting and quantified. Data represented as mean ± S.E.M. (n=3).

### Figure S1 (Related to Figure 1)

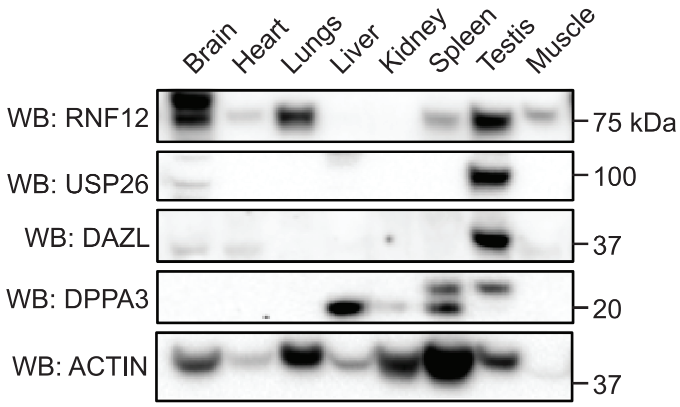

### Figure S2 (Related to Figure 4)

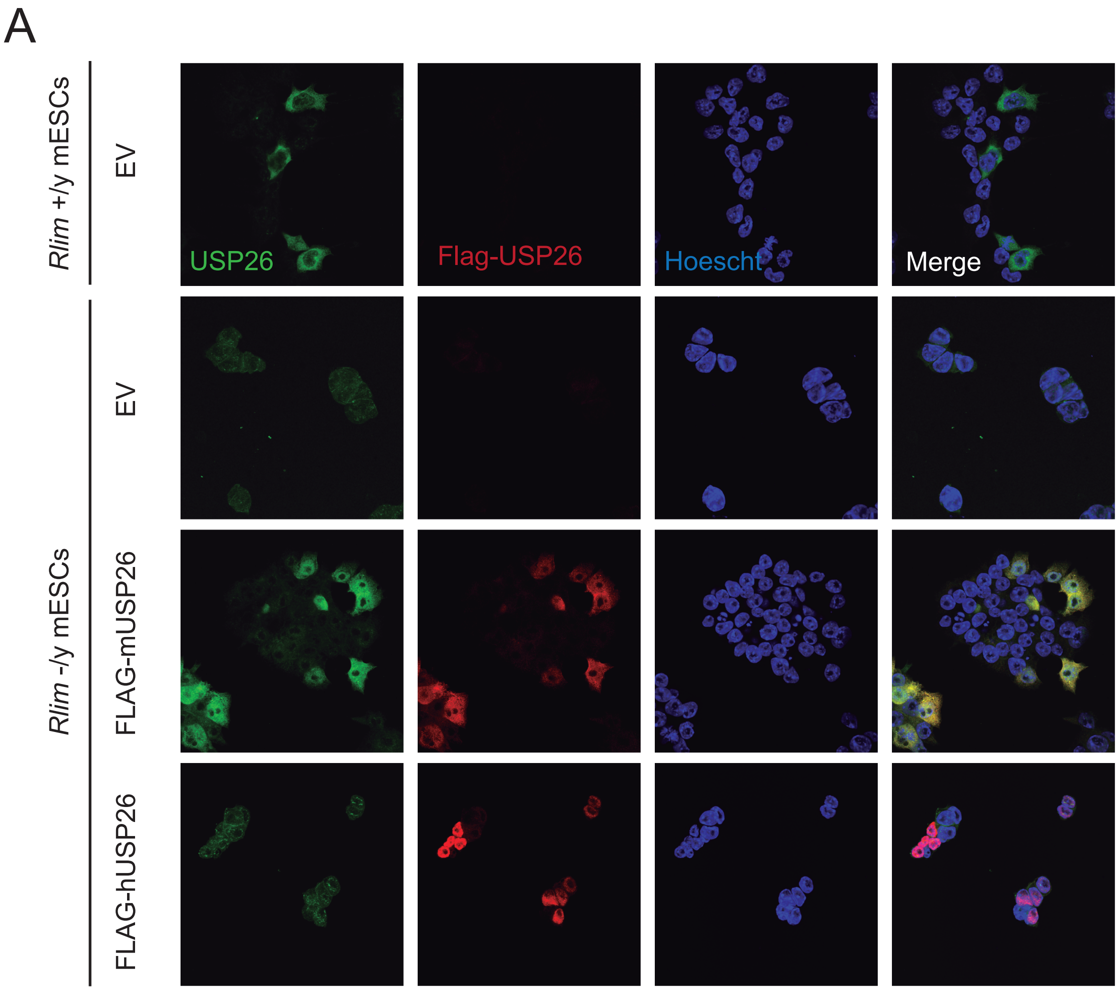


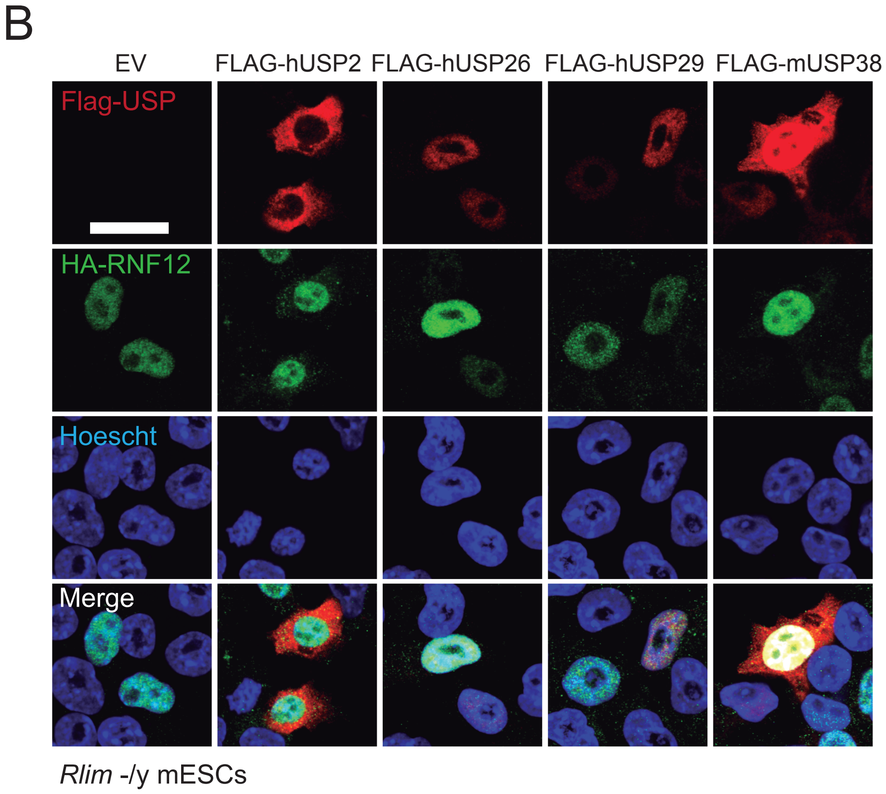

### Figure S3 (Related to Figure 5)

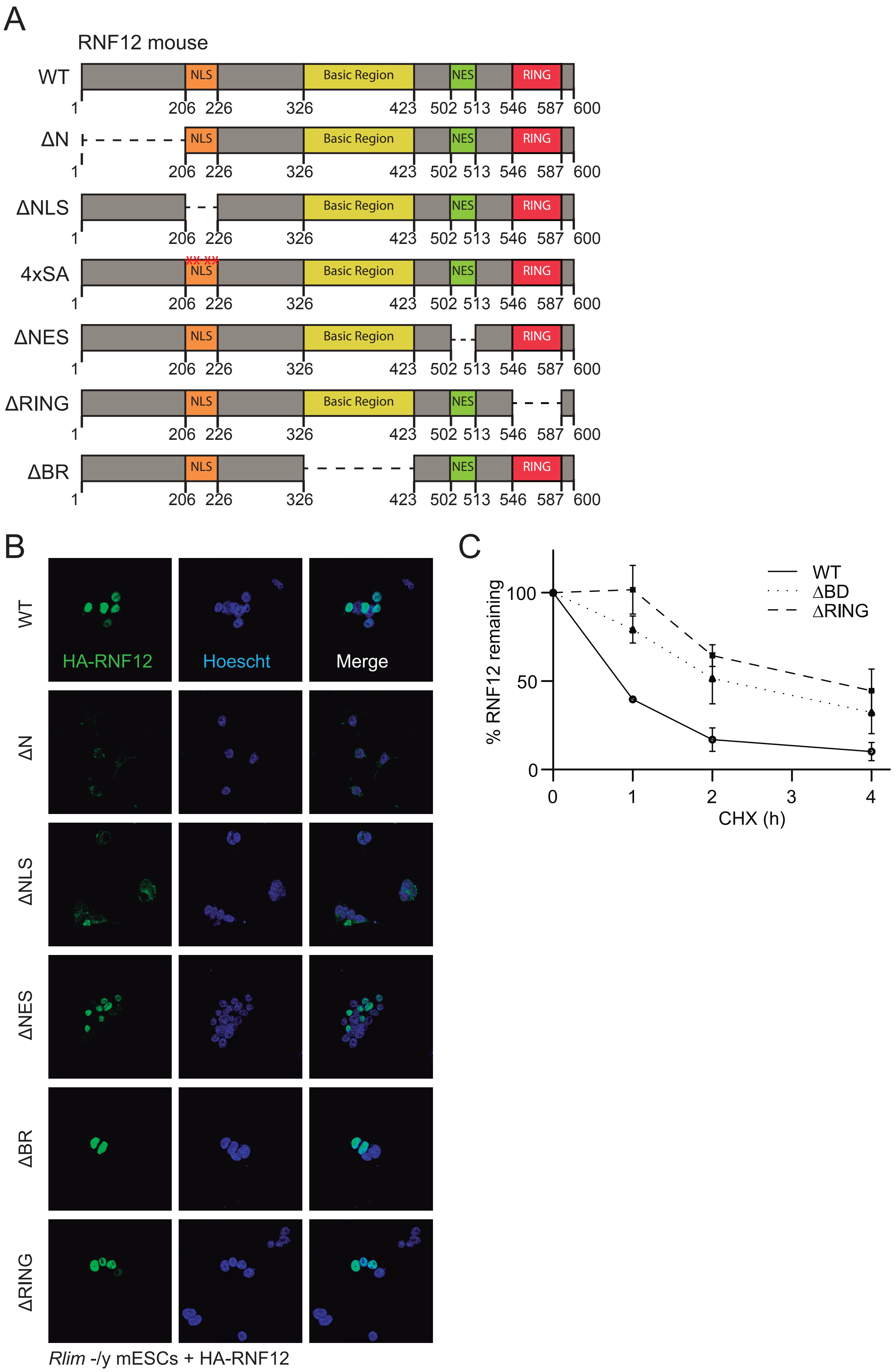
